# Trial-to-trial similarity and distinctness of muscle synergy activation coefficients increases during learning and with a higher level of movement proficiency

**DOI:** 10.1101/2023.09.19.558460

**Authors:** Paul Kaufmann, Willi Koller, Elias Wallnöfer, Basilio Goncalves, Arnold Baca, Hans Kainz

**Affiliations:** Department of Biomechanics, Kinesiology and Computer Science in Sport, Centre for Sport Science and University Sports, University of Vienna, Vienna, Austria; Neuromechanics Research Group, University of Vienna, Vienna, Austria

## Abstract

Muscle synergy analyses are used to increase our understanding of motor control. Spatially fixed synergy vectors coordinate multiple co-active muscles through activation commands, known as activation coefficients. To better understand motor learning, it is crucial to know how synergy recruitment varies during a learning task and different levels of movement proficiency. Within one session participants walked on a line, a beam, and learned to walk on a tightrope – tasks that represent different levels of proficiency. Muscle synergies were extracted over all conditions and the number of synergies was determined through the knee-point of the total variance accounted for (tVAF) curve. We found that the tVAF of one synergy decreased with task proficiency (line < beam < tightrope). Additionally, trial-to-trial similarity and distinctness of synergy activation coefficients increased with proficiency and after a learning process. We conclude that precise adjustment and refinement of synergy activation coefficients play a crucial role in motor learning.

## 1 Introduction

The underlying mechanisms, by which the central nervous system controls movements and adapts during learning new movements, are still not fully understood. One common theory in in the field of motor control implies the idea of muscle synergies [1–3]. Put simply, muscle synergies refer to groups of co-active muscles, termed synergy vectors or motor modules, which are recruited by activation coefficients, corresponding to time-dependent control inputs of the central nervous system [1, 3]. In line with Bernstein’s levels of movement construction [4, 5], this simplifies the complex coordination of the large number of muscles in the human body by controlling the activation of a limited number of spatially fixed, and temporally independent motor modules, rather than individually controlling each muscle.

Over the last two decades, muscle synergies, extracted from electromyography (EMG) recordings have been studied in healthy and pathological populations across various tasks. These studies have demonstrated the recruitment of similar motor modules in different movements, strengthening the concept of spatially fixed synergy vectors. So-called shared synergies describe similar movement fragments, which correspond to physical subtasks with the same mechanical goals [6]. For example, shared synergies were found between walking and cycling [7], walking and slipping [6], walking and standing reactive balance tasks [8], stepping and non-stepping postural behaviors [9], seated and standing cycling [10, 11] or overground and treadmill running [12]. To describe the complexity of motor control, the total variance in muscle activity accounted for (tVAF) by a given number of synergies, and the number of needed synergies (NoS) are widely utilized parameters. For instance, less synergies and higher tVAF – indicating lower motor complexity – were found in individuals with cerebral palsy [13, 14] or stroke [15, 16] compared to unimpaired populations, and in younger compared to older adults during walking [17].

It is generally accepted, that generating identical movements on successive attempts is impossible, due to an inherently noisy nervous system [18]. This noise can arise from either the central nervous system through movement planning or peripherical structures (i.e.: force production by muscles). In 2017, Dhawale et al. [19] reviewed recent studies regarding movement variability in motor learning, and concluded, that variability in the planning space is more likely a feature of motor system plasticity that drives motor learning, rather than unwanted noise. Moreover, this trial-to-trial variability decreases with increasing task proficiency [19–22], aligning with the principles of reinforcement learning [19]. Reinforcement learning theory suggests that a system learns new behaviors through trial-and-error [23]. Motor commands that lead to favorable outcomes (i.e.: successful execution of a movement task) are repeated, reinforced, and refined in subsequent attempts. In a study by Wu et al. [20] participants were trained to replicate a curve shape using hand trajectories in a reaching task. They found that individuals who displayed higher kinematic variability prior to training showed faster rates of learning. Hence it seems that variability during the learning process increases the likelihood of finding the optimal motor command.

To date, only few studies have examined the role of muscle synergies in movement learning. For instance, Sylos-Labini et al. [24] compared walking trial-to-trial variability of temporal synergy activations across different age groups, ranging from neonates to adults. They observed a decrease in variability during locomotor development. Consistent with a prior study on locomotor development [25], authors revealed that motor complexity and the number of synergies increased with age. In adults, changes of activation coefficients variability correlated with changes in bowling scores across sessions [26]. Comparing professional ballet dancers with individuals without any dancing or gymnastics experience, Sawers et al. [27] revealed higher trial-to-trial similarity with higher beam walking proficiency. Additionally, dancers showed lower variability within synergy vectors and higher spatial distinctness between synergy vectors. Similarly, dance-based rehabilitation in individuals with Parkinson’s disease improved the consistency and distinctness of synergy vectors [28]. All the mentioned studies were limited by either inter-participant variability [24, 29–32], or inter-session variability [31–33], which can be attributed to individual motor control differences and variations in skin-electrode impedance and electrode position.

To the best of our knowledge, no study has yet examined changes in muscle synergies using a within-participant, within-session protocol. Therefore, the present study addresses this research gap. Briefly, each participant walked on a line, a beam, and a tightrope. The choice of these three tasks was based on the progression from an easy, daily task with highest movement proficiency (line) to a more uncommon task that was still manageable for participants (beam) and finally to a new task, which could be learned within one data collection session (tightrope). The twofold aim of the study was to examine if motor complexity, trial-to-trial similarity of activation coefficient and activation coefficient distinctness differs: (1) between an early and a late stage of a learning process (i.e.: learning to walk on a tightrope); (2) between common and less common movement tasks – addressing movement proficiency. Subsequently, we investigated, if the contribution of synergies changes among learning or proficiency changes. The study primarily focused on muscle synergies, but trial-to-trial similarity of EMG envelopes and joint angles were also analyzed to gain a comprehensive understanding of variability in motor learning. Additionally, the study investigated whether the amount of muscle activity changes after learning, building on previous findings by Donath et al. [34], who showed decreased muscle activity after slackline training. We hypothesize that motor complexity, activation coefficient distinctness and trial-to-trial similarity of synergy activation, EMG envelopes, and joint angles (1) gets higher during learning, and (2) is higher in more common movements. Furthermore, the study hypothesizes that the amount of muscle activity decreases during learning (1) and is lower in more common movements (2).

## 2 Materials and Methods

### 2.1 Participants

This study involved ten healthy participants (age: 25.2 ± 3.34 years; bodyweight: 69.9 ± 7.34 kg; height: 1.76 ± 0.09 m; body-mass-index: 22.63 ± 1.51; 6 men and 4 women) without neurological or orthopedic impairments, who were not able to walk on a slackline or tightrope beforehand. The study was approved by the ethics committee of the University of Vienna (reference number: 00820) and participants gave written informed consent.

### 2.2 Experimental Setup and Data Collection

Each participant walked under different tasks: (a) a line taped on the ground (LINE; length: 310 cm; width: 1.4 cm); (b) a beam (BEAM; length: 341.5 cm; width: 10 cm; height: 28.5cm) and (c) a tightrope (TIGHTROPE; length: 363 cm; diameter: 0.9 cm; height: 363 cm) spanned between two platforms (Figure 1). The learning process for walking on the TIGHTROPE was divided into two stages: TRfail and TRsucc. TRfail included the first five attempts where participants were able to perform at least one full gait-cycle of the right leg but were not able to successfully balance over the entire tightrope. TRsucc included the attempts where participants successfully balanced over the tightrope in four out of five consecutive attempts. A successful attempt was defined as walking over the whole tightrope and maintaining balance on the second platform. If a participant was able to successfully balance over the TIGHTROPE in two out of the first five attempts, the difficulty of the task was increased with visual constraints, by either an eye-patch over the left eye, or further by closing both eyes (if two of the first five trials were successful with the eye-patch). The conditions were recorded in the following order: (1) LINE-walking (startLine), (2) BEAM-walking (startBEAM), (3) start of learning process on the TIGHTROPE (TRfail) until (4) the end of learning process (TRsucc), (5) BEAM-walking (endBEAM), and (6) Line-walking (endLINE). To ensure consistent visual constraints across tasks for later comparisons, five trials with opened eyes, an eye-patch over the left eye and both eyes closed were recorded each time (start and end) for LINE and BEAM. As data from the first right stance phase was further analyzed, we aimed to minimize transient accelerations at the onset step [14, 17, 35], by instructing participants to start each trial with their left leg. Only the stance phase was analyzed, to neglect highly variable movement times between stance and swing phases across conditions [32, 36]. No additional constraints for pause time, step cadence, step length, or hints for walking over the TIGHTROPE were given, to provide self-directed learning.

**Figure 1:**
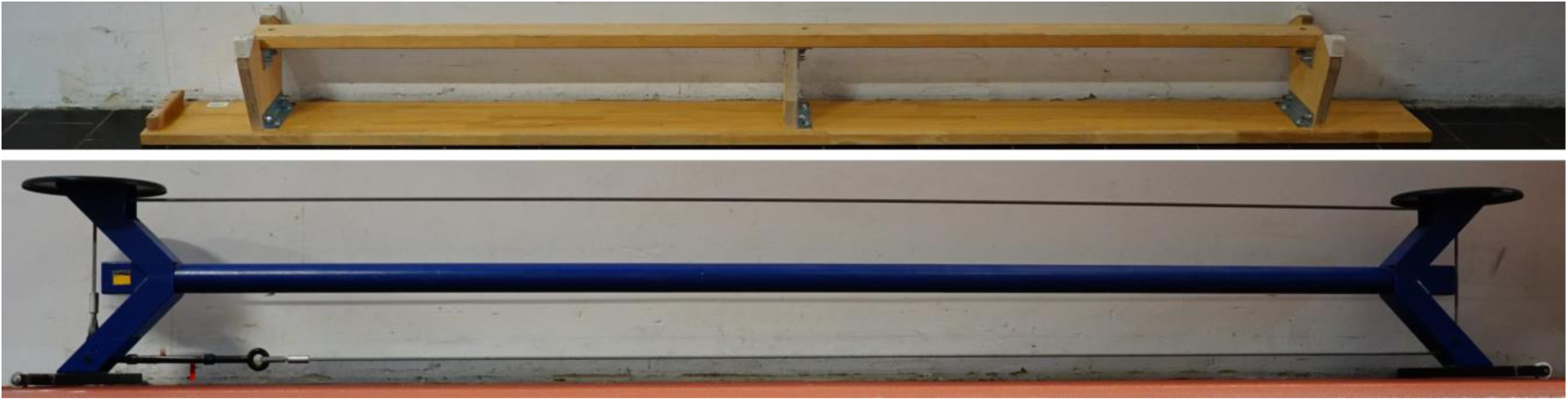
Top image shows the upside-down gymnastics bench which was used for BEAM conditions. Bottom image shows the THIGHTROPE mounted on a rack between two platforms.

Prior to the data collection, thirteen surface EMG sensors (eleven PicoEMG and two Mini Wave Infinity, Wave Plus wireless EMG system, Cometa, Milan, Italy) were placed on the trunk and right limb following the Seniam guidelines (Seniam.org) and recommendations from previous studies [37–39]: tibialis anterior (tib_abt), peroneus longus (per_long), soleus, gastrocnemius medialis (gast_med), vastus lateralis (vast_lat), rectus femoris (rect_fem), biceps femoris (bic_fem), semitendinosus (sem_tend), gluteus maximus (glut_max), rectus abdominis (rect_abd), extensor obliques (ext_obli), multifidus (multifid) and erector spinae iliocostalis (erec_spin). A baseline EMG signal of several seconds was collected (EMG_base) while participants lied in a supine and relaxed position on a massage table. The standard Vicon Plug-in-Gait marker set (Vicon, Oxford, UK), including 21 reflective markers, were placed on the legs and the trunk of each participant [40]. The heel and toe markers were placed on the shoes of participants, similar to Paterson et al. [41]. A 12-camera 3D motion capture system (Vicon, Oxford, UK) was used to record marker trajectories with a sampling rate of 200 Hz, EMG data with 1000 Hz and ground reaction forces of one force plate with 1000 Hz (Kistler Instrumente, Winterthur, Switzerland), simultaneously. In addition, participants wore in-shoe force sensor soles (loadsol^®^, Novel, Munich, Germany), which were used to determine stance phases. Insoles data was captured with 200 Hz (loadsol-s android application version 1.7.63) on a mobile phone (Huawei P30 Lite, Huawei, Shenzhen, China) and brought to zero level every 5 to 10 trials to minimize errors due to sensor drifts. Foot contact instances were determined by vertical contact forces over 30 Newton via custom scripts. Time synchronization between the Insole and Vicon data was achieved by participants stepping on a force plate at the beginning of each trial. The experimental data was captured and processed using Vicon Nexus 2.12 software (Vicon, Oxford, UK). Subsequent analyses were conducted using Gnu Octave version 6.2.0 [42] and MATLAB (R2022a, Mathworks Inc., Natick, USA).

### 2.3 EMG processing

Raw EMG signals were high-pass filtered at 25 Hz (4^th^-order Butterworth zero lag filter) to remove movement artefacts [14, 43–45], demeaned, full-wave rectified and low-pass filtered at 7 Hz (4^th^-order Butterworth zero lag filter), similar to previous gait studies [46–50]. The low-pass cutoff frequency of 7 Hz was chosen as a compromise between the different movement times (supplementary Figure 1). After filtering, baseline noise was removed by subtracting the root-mean-square of the filtered EMG_base signal, to improve signal-to-noise ratio [51–54], and resulted negative values were set to zero. Based on a visual inspection of raw and filtered EMG envelopes, trials with artefacts were removed, resulting in four to five remaining trials per condition. Afterwards signals were time-normalized to 101 data points (100% of stance phase) and amplitude normalized to values between 0 and 1, where an amplitude of 1 was equal to the maximum activation amplitude of a muscle among all trials [16, 35, 36, 48, 55].

### 2.4 Synergy extraction and determining the number of synergies

For each participant, processed EMG signals of trials from all conditions were concatenated [10, 56] and muscle synergies were extracted according to the spatial/synchronous synergy model. According to this model, motor control of muscle activations (EMG signals), is described by a linear combination of a fixed spatial synergy vector W and a time-varying activation coefficient C [4, 36, 52]. Non-negative-matrix-factorization (NNMF) has been shown to be the most appropriate method for extracting muscle synergies in walking [50]. Therefore, we used the “nmf_bpas” octave function [57], an advanced algorithm of the classic NNMF [58–60] to extract one to twelve (number of muscles −1) muscle synergies. Instead of random inputs, the NNMF was initialized with outputs of the nonnegative single-value-decompensation with low-rank correction algorithm [61] to improve NNMF [52, 61–63]. Extracted synergy vectors were normalized to 1 based on their maximum values, and activation coefficients were multiplied by the same normalization values, to keep their product constant [64, 65]. More information regarding the synergy extracting procedure is provided in the supplementary material.

The total variance accounted for (tVAF) was calculated for each number of extracted synergies (1 to 12). It quantifies the reconstruction accuracy after the factorization, and is defined as the uncentered Pearson’s correlation in percentage [49]. To determine the number of synergies that represents motor control across all conditions (NoS), knee point analysis was employed [36, 44, 49, 52, 66]. The knee-point (v) was defined as the point on the tVAF curve that exhibits the smallest angle among three adjacent points (v-1, v, v+1). This approach assumes that beyond the knee-point, only unstructured data or noise is explained by additional motor modules [66]. It was preferred over threshold-based methods, as it has been shown to perform better [49] and is not affected by different low-pass filter cutoff frequencies [44]. We further constrained our analysis by exclusively determining the knee-point for synergies with a tVAF exceeding 95%. This widely used threshold [46, 49, 54, 56, 67–69] was added based on visually observing sharp jumps in some tVAF curves, likely caused by the split of a synergy due to salient features [70].

### 2.5 Assessment of trial-to-trial similarity

The trial-to-trial similarity of synergy activation coefficients, EMG envelopes and joint angles were all quantified based on the same three parameters: the Pearson correlation coefficient (r), the maximum value of the normalized cross-correlation coefficient (r_max_) and the lag time (lag% in % of the stance phase) where r_max_ occurred which represents the time shift between two curves. These parameters are widely used to quantify variabilities in synergy, EMG and kinematic waveforms [7, 10, 11, 64, 71–73]. We calculated them for every pairwise combination of trials in each condition within each synergy, muscle, and joint. The averaged value per condition represents the overall trial-to-trial similarity for synergy activation coefficients, EMG envelopes and joint angles.

### 2.6 Synergy analyses

We computed the tVAF using the EMG signals, synergy vectors and activation coefficients of each condition. Then, tVAF of one synergy (tVAF1) and tVAF at NoS (tVAFNoS) were compared across conditions to evaluate movement complexity (tVAF1) and the goodness of reconstruction (tVAFNoS). The distinctness of activation coefficients was determined by calculating the average value of all pairwise combinations of activation coefficients from different synergies within each trial for each condition. High values of r and r_max_, along with small time-shifts (lag%), indicate a similarity in timing and a substantial amount of overlapping in synergy activations [16, 62].

Additional to the overall trial-to-trial similarity of each condition, we aimed to reveal, if differences in the variability just occur in some synergies. To classify similar synergy vectors among participants, we used k-means clustering (kmeans function in Octave – see supplementary material) similar to recent synergy studies [24, 26, 30, 48, 74]. We computed the k-means clustering solution for a range of two to twelve clusters and repeated the process 100 times. For each repetition and each number of clusters, we calculated the average silhouette value [75]. The optimal number of clusters was then determined on the point at which the maximum silhouette values plateaued – indicating small within- and high between-cluster distances [26] (Figure 4). Trial-to-trial similarity parameters (r, r_max_, lag%) were calculated for synergies within the same cluster, for each condition. For instance, if a cluster consisted of eight synergy vectors, the trial-to-trial similarity of that cluster was determined by averaging the trial-to-trial similarity values of the eight synergies. To examine the task-specific relevance of individual synergies, tVAF by each synergy was computed for every trial. These tVAF values were then averaged across synergies within the same cluster.

### 2.7 EMG analyses

To quantify changes in the amount of muscle activity, the root-mean-square (RMS) of the preprocessed EMG signals of every trial was calculated and averaged across trials of the same condition, within each muscle. Additionally, to the overall trial-to-trial similarity (section 2.5), correlation values were also averaged for each muscle to evaluate, if variability in activation patterns only occurred in some muscles (results provided in supplementary material).

### 2.8 Kinematic analyses

Joint angles were computed with OpenSim [76] using the recently introduced addBiomechanics.org application [77]. This application uses a bilevel optimization and enables to personalize musculoskeletal models and calculate joint angles in an easy and efficient way. We used the default option with the Rajagopal2015 model for human gait [78]. The computed joint angles were smoothened using a 6 Hz low-pass filter (4^th^-order Butterworth zero lag filter) and time normalized to 101 datapoints of the stance phases. The following joint angles of the right leg and trunk were examined: ankle plantar-/dorsiflexion, knee flexion/extension, hip flexion/extension, hip ab-/adduction, hip internal/external rotation, lumbar flexion/extension, lumbar medial/lateral bending, and lumbar internal/external rotation. In addition to the overall trial-to-trial similarity (section 2.5), correlation values were also averaged for each joint separately, to evaluate, if variability in kinematics only occurred in some joints (results provided in supplementary material).

### 2.9 Statistics

We employed a two-way repeated measures ANOVA with TASK (LINE, BEAM, TIGHTROPE) and TIME as factors on all variables described above. The first time point (START) consisted of startLINE, startBEAM, and TRfail, while the second time point (END) included endLINE, endBEAM, and TRsucc. TASK was used to assess differences regarding task commonness – our second research question-including post hoc pairwise comparisons with Bonferroni correction. To address our first research question, i.e. changes during the learning process, – we calculated contrasts between TRfail and TRsucc. Furthermore, contrasts between startLINE and endLINE were examined as a baseline to assess the stability of the analyzed variable, as no differences were anticipated between the two LINE conditions. Additionally, contrasts between startBEAM and endBEAM were analyzed to explore potential transfer effects of learning from one balancing task (TIGHTROPE) to another (BEAM). Contrasts were conducted only if a significant difference was observed in any of the ANOVA outcomes (TASK, TIME, TASK*TIME). Prior, sphericity was checked with Mauchly-test (if necessary, Greenhouse-Geisser correction was applied), and normal distribution was verified with Shapiro Wilk-test. If the requirement of normal distribution was violated, an aligned-rank-transformation was performed. This transformation enabled us to conduct factorial ANOVA’s on nonparametric data [79–81] and was utilized with ARTool 2.1.2 software (Washington, USA). Statistical analyses were performed with JASP 0.17.2 (Amsterdam, Netherlands). The alpha level was set at 0.05, and the results were reported at three levels of significance: p < 0.05, p < 0.01, and p < 0.001.

## 3 Results

Participants required 2.6 ± 1.4 attempts (range: 1 to 5) to perform their first complete gait-cycle with the right leg (TRfail) and 49.4 ± 22.8 attempts (range: 12 to 101) to complete the learning task (TRsucc). Two participants walked on the TIGHTROPE with visual constraints (1x eye-patch, 1x closed eyes).

### 3.1 Muscle synergy analyses

An average of 5.9 ± 1.1 NoS was determined among participants. For tVAF1 a significant effect of TASK (p < 0.001) was observed. Post hoc comparisons revealed that tVAF1 was higher in BEAM compared to LINE (p < 0.01) and TIGHTROPE was higher than both LINE and BEAM (p < 0.001). There were no significant differences in tVAFNoS. Regarding the distinctness of activation coefficients, the ANOVA revealed a significant effect of TASK for r (p < 0.05), where activation coefficients were more correlated to each other (p < 0.05) in TIGHTROPE compared to LINE and BEAM. There was no significant difference for r_max_, but a significant effect of TASK (p < 0.001) and TIME (p < 0.05) in lag%. The lag% was higher in LINE than BEAM (p < 0.01) and TIGHTROPE had the lowest %lag (p < 0.001) (Figure 2).

**Figure 2:**
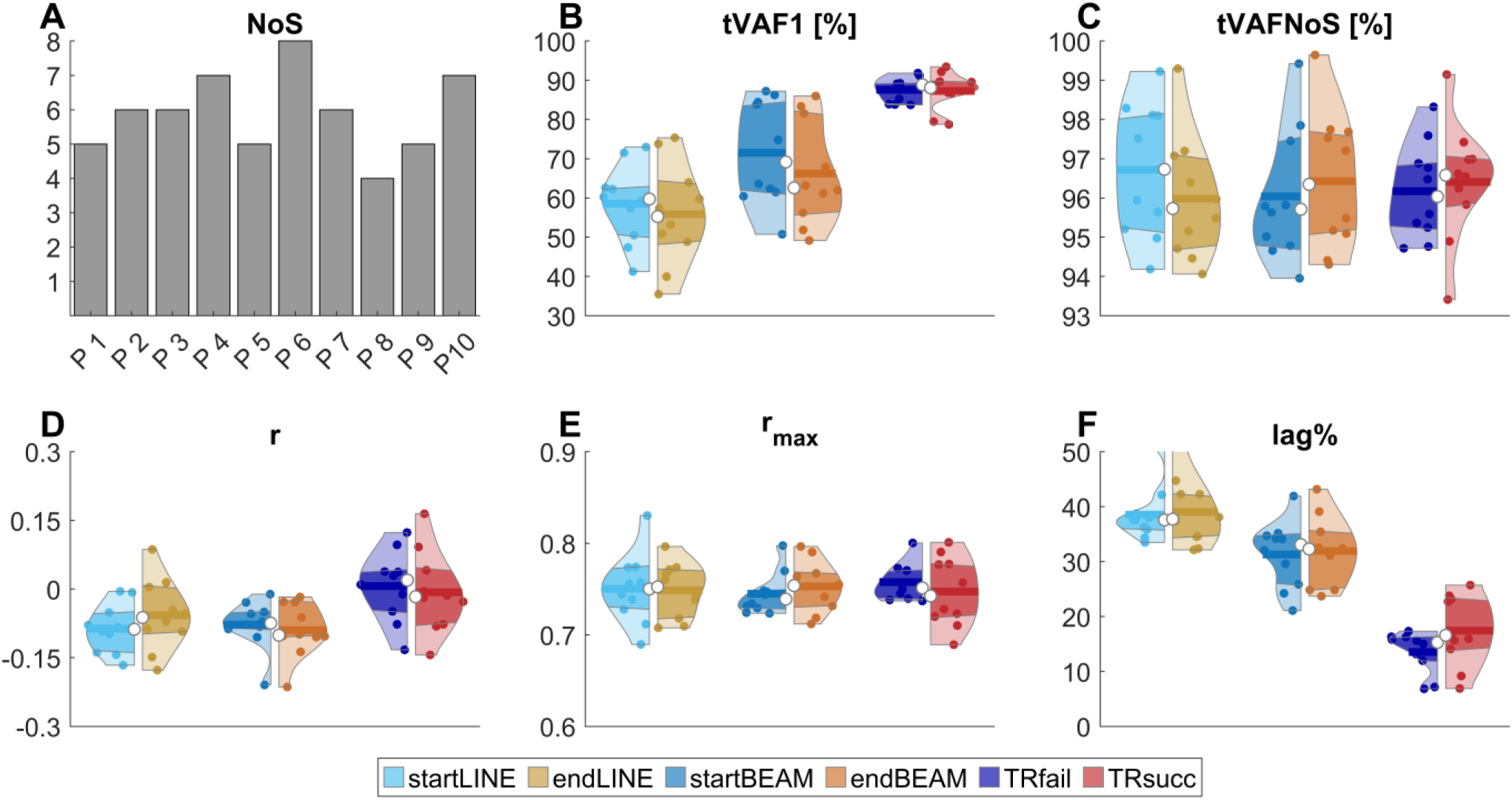
**A**: bars show the number of required synergies (NoS) for each participant (P1 – P10). **B-C**: the total variance accounted for one synergy (**B**: tVAF1) and NoS (**C**: tVAFNoS). **D-F**: Synergy activation coefficient distinctness measured by Pearson correlation (**D**: r), maximum cross-correlation coefficient (**E**: r_max_) and lag at r_max_ (**F**: lag%). Violin plots: each colored circle represents one participant; thick lines represent mean values; white circles indicate median values; dark areas indicate quartiles.

Trial-to-trial similarity measured by r and r_max_ was affected by TASK (p < 0.001) and TIME (p < 0.01). r and r_max_ was the highest in LINE, followed by BEAM (r_max_: p < 0.01; r < 0.001) and lowest in TIGHTROPE (both: p < 0.001). Contrasts showed an increase in similarity from startBEAM to endBEAM (both: p < 0.05) and TRfail to TRsucc (r_max_: p < 0.05; r: p < 0.001). There was no difference in lag% (Figure 3).

**Figure 3:**
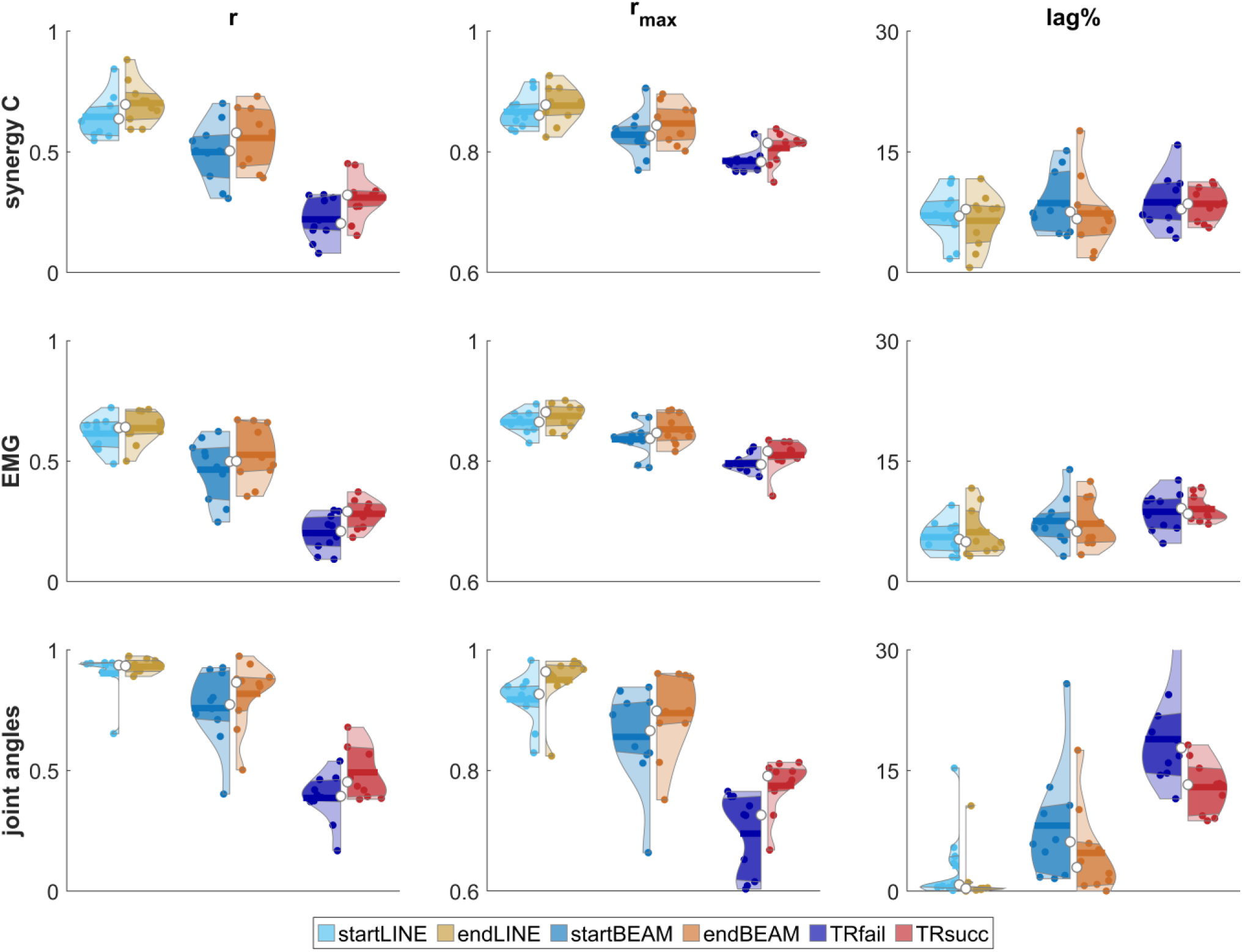
Overall trial-to-trial similarity of synergy activation coefficients (C, top row), electromyography envelopes (EMG, middle row) and joint angles (bottom row), measured by Pearson correlation (r), maximum cross-correlation coefficient (r_max_) and lag at r_max_ (lag%). Violin plots: each colored circle represents one participant; thick lines represent mean values; white circles indicate median values; dark areas indicate quartiles.

Silhouette analyses yielded six clusters (Figure 4) which are indicated by # in the following paragraphs. Low tVAF values indicate low contribution of synergies to the condition. The tVAF of all clusters was significantly affected by TASK (#5: p < 0.05; #1, 3: p < 0.01; others: p < 0.001). In #4, tVAF of BEAM was lower than LINE (p < 0.05) and the lowest in TIGHTROPE (p < 0.001). For the other clusters, tVAF of TIGHTROPE was higher than BEAM (#5, 6: p < 0.05; #2: p < 0.001) and LINE (#1, 3, 5: p < 0.01; #2, 6: p < 0.001). In BEAM it was higher than LINE (#2: p < 0.01). For #2, ANOVA also revealed a significant effect of TIME (p < 0.05), with lower tVAF in START compared to END, and the interaction TASK × TIME (p < 0.01). In #6, contrasts showed a decrease of tVAF over time in BEAM (p < 0.05).

**Figure 4:**
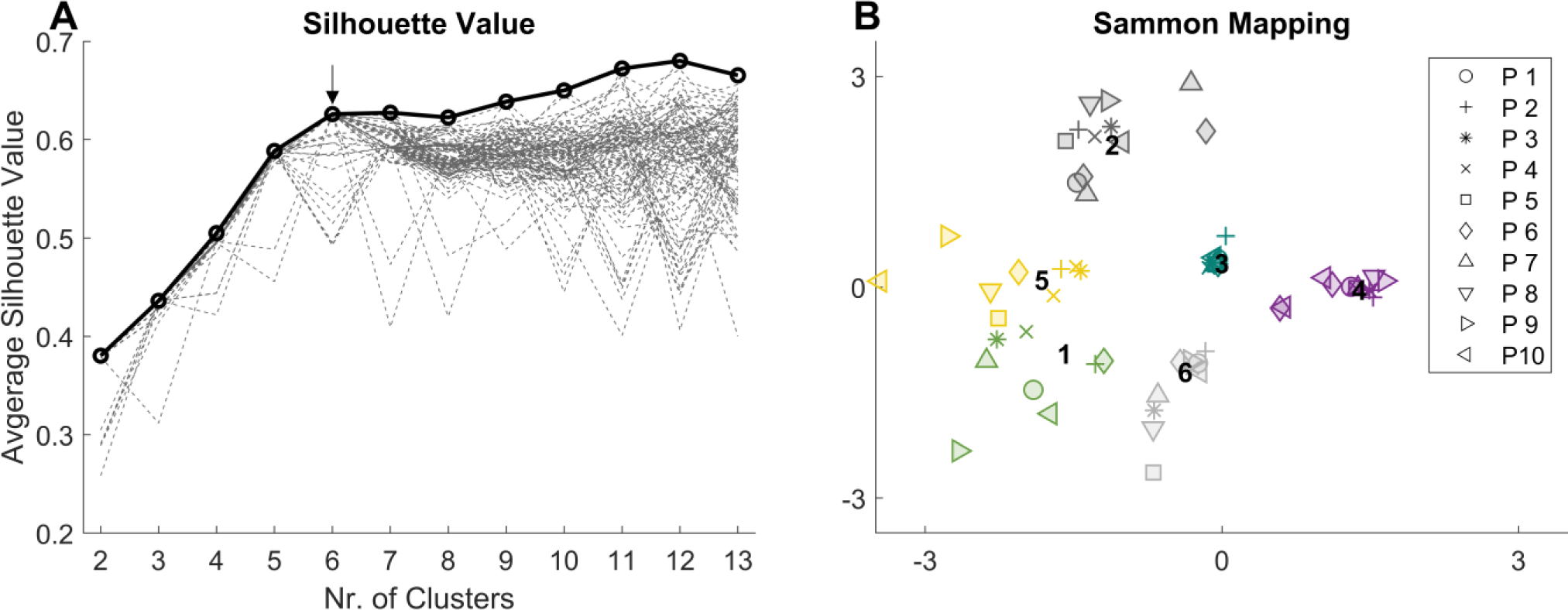
**A**: dashed lines show the average silhouette value for each clustering repetition (1 to 100). The arrow indicates the number of clusters, at which the maximum of averaged silhouette values among repetitions (solid line/circles) plateaued. **B**: sammon mapping [93] of the six clusters. Marker-styles indicate different participants (P1 – P10), and marker-colors indicate different clusters. Numbers (1 to 6) indicate the position of the clusters’ centroids.

Trial-to-trial similarity of cluster 1, 2, 4 and 5 was significantly affected by TASK in r and r_max_ (r_max_ #5: p < 0.05; r #5: p < 0.01 others: p < 0.001), with higher LINE than TIGHTROPE for r (p < 0.001) and r_max_ (#5: p < 0.05; others: p < 0.001) and higher BEAM than TIGHTROPE for r (#5 p < 0.01; others: p < 0.001). r_max_ was higher in BEAM than TIGHTROPE in cluster 1 (p < 0.01), 2 and 4 (p < 0.001). Correlation was higher in LINE than BEAM in cluster 1, 2 (r and r_max_: p < 0.05) and 4 (r: p < 0.01; r_max_: p < 0.001). The lag% revealed a significant effect of TASK for cluster 1, 3, 4 and 5(#3: p < 0.05; #5: p < 0.05; #1, 4: p < 0.001). LINE had lower lag% than BEAM (#1: p < 0.05) and TIGHTROPE (#5: p < 0.01; #1, 4: p < 0.001). BEAM had lower lag% compared to TIGHTROPE (#1, 5: p < 0.01). Contrary, #3 had the lowest lag% in TIGHTROPE compared to the other two conditions (p < 0.05).

Significant effects of TIME were found for r in cluster 1 (p < 0.05), for r_max_ in cluster 4 and 6 (p < 0.05) and for lag% in cluster 4 (p < 0.05) with lower correlations and higher lag% in START compared to END. A significant effect of TASK × TIME was only found for r in cluster 4 (p < 0.05). Contrasts revealed a significant increase of r or r_max_ from startLINE to endLINE in cluster 6 (r_max_: p < 0.05), from startBEAM to endBEAM in cluster 2 (r: p < 0.01) and from TRfail to TRsucc in cluster 1 (r: p < 0.05).

### 3.2 EMG analysis

Overall trial-to-trial similarity of EMG envelopes measured by r and r_max_ was significantly affected by TASK (p < 0.001), with LINE showing the highest correlation, followed by BEAM, and TIGHTROPE at last (r LINE vs BEAM: p < 0.01; others: p < 0.001). TIME influenced r (p < 0.01) and r_max_ (p < 0.05) and contrasts revealed lower r and r_max_ (p < 0.05) for startBEAM compared to endBEAM, and an increase in r (p < 0.01) between TRfail and TRsucc. The lag% was significantly affected by TASK (p < 0.01), with higher values in TIGHTROPE compared to LINE (p < 0.01). (Figure 3, Figure 7).

The amount of muscle activation measured by RMS revealed a significant effect of TASK, in all muscles, apart from soleus (glut_max: p < 0.01; others: p < 0.001). RMS of gast_med was lower in TIGHTROPE than BEAM (p < 0.05) and LINE (p < 0.001). For the other muscles, RMS was higher in TIGHTROPE than BEAM (glut_max: p < 0.05; tib_ant, bic_fem: p < 0.01; others: p < 0.001) and LINE (p < 0.001). In four muscles BEAM was also higher than LINE (rect_fem, multifid: p < 0.05; per_long, erec_spin: p < 0.01). There was a significant effect of TIME (rect_fem; bic_fem, glut_max: p < 0.05; tib_ant soleus, gast_med, sem_tend, erec_spin: p < 0.01; vast_lat, rec_abd, ext_obli: p < 0.001), and TASK × TIME (ext_obli: p < 0.05; tib_ant, vast_lat, sem_tend: p < 0.01; rec_abd, multifid, erec_spin: p < 0.001) on muscle activations. Contrasts revealed a higher muscle activation in startLINE than endLINE for two muscles (gast_med: p < 0.05, sem_tend: p < 0.01), startBEAM than endBEAM for four muscles (soleus, rect_fem, rec_abd: p < 0.05; tib_ant: p < 0.01), and TRfail than TRsucc for ten muscles (tib_ant, soleus, gast_med, erec_spin: p < 0.01; vast_lat, sem_tend, glut_max, rec_abd, ext_obli, multifid: p < 0.001) (Table 1).

**Table 1:** Muscle activations (root-mean-square) for all conditions and muscles. M and SD represent the mean and standard deviation values across all participants. ANOVA revealed significant effects of TASK in all muscles apart from soleus. Significant differences observed by contrasts are indicated by *.

|  | LINE |  |  |  | BEAM |  |  |  | TIGHTROPE |  |  |  |
| --- | --- | --- | --- | --- | --- | --- | --- | --- | --- | --- | --- | --- |
|  | start |  | end |  | start |  | end |  | fail |  | succ |  |
|  | M | SD | M | SD | M | SD | M | SD | M | SD | M | SD |
| tib_ant | 0.13 | 0.07 | 0.14 | 0.07 | 0.22* | 0.09 | 0.16* | 0.07 | 0.33* | 0.07 | 0.27* | 0.08 |
| per_long | 0.19 | 0.05 | 0.17 | 0.05 | 0.24 | 0.04 | 0.23 | 0.04 | 0.37 | 0.06 | 0.35 | 0.07 |
| soleus | 0.26 | 0.07 | 0.23 | 0.07 | 0.27* | 0.06 | 0.24* | 0.04 | 0.31* | 0.10 | 0.25* | 0.08 |
| gast_med | 0.32* | 0.06 | 0.29* | 0.06 | 0.29 | 0.05 | 0.27 | 0.05 | 0.24* | 0.08 | 0.20* | 0.06 |
| vast_lat | 0.14 | 0.09 | 0.12 | 0.07 | 0.17 | 0.10 | 0.15 | 0.08 | 0.32* | 0.07 | 0.24* | 0.08 |
| rect_fem | 0.05 | 0.03 | 0.04 | 0.02 | 0.07* | 0.04 | 0.06* | 0.02 | 0.21 | 0.09 | 0.17 | 0.08 |
| bic_fem | 0.09 | 0.05 | 0.08 | 0.07 | 0.11 | 0.07 | 0.11 | 0.08 | 0.25 | 0.08 | 0.20 | 0.09 |
| sem_tend | 0.17* | 0.09 | 0.13* | 0.08 | 0.17 | 0.08 | 0.16 | 0.09 | 0.29* | 0.08 | 0.20* | 0.08 |
| glut_max | 0.12 | 0.05 | 0.11 | 0.04 | 0.15 | 0.06 | 0.15 | 0.05 | 0.22* | 0.09 | 0.18* | 0.08 |
| rec_abd | 0.03 | 0.03 | 0.02 | 0.03 | 0.05* | 0.05 | 0.04* | 0.03 | 0.17* | 0.08 | 0.10* | 0.08 |
| ext_obli | 0.05 | 0.03 | 0.04 | 0.02 | 0.09 | 0.06 | 0.07 | 0.03 | 0.23* | 0.07 | 0.17* | 0.07 |
| multifid | 0.12 | 0.04 | 0.14 | 0.06 | 0.17 | 0.04 | 0.16 | 0.07 | 0.28* | 0.04 | 0.21* | 0.06 |
| erec_spin | 0.05 | 0.02 | 0.05 | 0.03 | 0.10 | 0.04 | 0.08 | 0.05 | 0.29* | 0.03 | 0.20* | 0.05 |

### 3.3 Kinematic analysis

Overall trial-to-trial similarity of joint angles, quantified by r, r_max_ and lag%, was significantly affected by TASK (p < 0.001). LINE exhibited the highest correlations and lowest lag%, followed by BEAM, and TIGHTROPE (r_max_ LINE vs BEAM: p < 0.01; lag% LINE vs BEAM: p < 0.05; others: p < 0.001). There was a significant effect of TIME on r (p < 0.05), with lower r in START compared to END, and a significant interaction effect of TASK × TIME (p < 0.05). For r_max_, TIME had a significant effect (p < 0.01), with an increase observed between START and END. All contrasts were significant (p < 0.05). Likewise, lag% was significantly influenced by TIME (p < 0.01). Contrasts revealed higher lag% in startLINE and TRfail compared to endLINE and TRsucc, respectively (p < 0.05) (Figure 3).

## 4 Discussion

The aim of the study was to increase our insights in motor learning using synergy analysis by employing a within-participant, within-session study design. We observed higher distinctness and trial-to-trial similarity of activation coefficients with increasing movement proficiency. Furthermore, the analyses revealed that the contribution of specific synergies varies across tasks, and muscle activity decrease throughout the learning process.

Over half a century ago Bernstein [5] proposed, that people restrict the number of degrees of freedom to simplify coordination in early learning stages. Steele et al. [67] found higher overlapping of synergy activation coefficients with the occurrence of biomechanical and task constraints. The current study showed higher tVAF1 and overlapping of synergy recruitment – both indicating higher coactivation of synergy vectors – in movements with lower proficiency. Taken these findings together, we suggest that freezing the number of degrees of freedom in early learning is a result of coactivating synergy vectors. In consequence, high tVAF values might be caused by overlapping synergy activations and not necessarily mean a simpler motor control due to a decreased number of synergies. This theory is supported by previous studies on impaired and unimpaired populations. Clark et al. [16] found similar synergy vectors in locomotion for stroke survivors and unimpaired individuals, if the same number of synergies were extracted, rather than the number determined by a tVAF threshold. The authors concluded that not the spatially synergy vectors differ, but they were computationally merged through the factorization algorithm due to their overlapping recruitment profiles. Similarly, merging of synergy vectors was found in locomotion of individuals post-stroke [82] and with Parkinson’s disease [36], and in reaching tasks after cortical lesions [83]. A higher amount of shared synergies between overground walking and balancing tasks was found in expert dancers compared to individuals with no dancing experience [8, 27], in post-stroke survivors compared to unimpaired individuals [84], and after a dance-based rehabilitation in individuals with Parkinson’s disease [28]. Two of these studies [27, 28] also found lower distinctness of synergy vectors in groups with fewer shared synergies. The lower distinctness of computed vectors may be a result of higher overlapping of activation coefficients, which can compromise the accuracy of extracted synergy vectors. This phenomenon has been observed in previous studies on real and simulated datasets [62, 66, 67], where increased temporal overlap of activation coefficients led to merging of synergies due to the underlying assumptions of factorization algorithms. Consequently, these inaccurately extracted synergy vectors could explain the lower number of shared synergies. In the current study we also found a low number of shared synergies when computing them separately for each condition, but similar synergies when computing them over all conditions (see supplementary material). This suggests that with proficiency overlapping of activation coefficients reduced, rather than the number of shared synergies changed. This concept should be addressed in further studies.

An important feature of motor learning is motion fusion, also called coarticulation, which describes the combination of individual movement primitives into a smooth action. More precisely, the velocity peaks of two movements gradually disappear during learning. Typically, motion fusion is assessed by examining velocity peaks in hand trajectories during tasks that involve precise movements, such as following a specific curvature on a monitor. [85–88]. At first glance, our findings of higher activation distinctness with proficiency may seem to contradict the concept of motion fusion. However, further analysis (results not presented) revealed that the timing of velocity peaks in knee and ankle flexion/extension became more synchronized with higher proficiency. This suggests that improved coordination of synergy activation timing leads to motion fusion and ultimately results in smoother movements. Even thought we did not find significant changes in tVAF1 and distinctness between TRfail and TRsucc, these factors might change during learning and were potentially not significantly affected in the current study due to still quite low movement proficiency (4 out of 5 successful attempts) after learning.

Analysis on muscle activations revealed that all muscles apart of the gastrocnemius medialis were more activated in TIGHTROPE compared to LINE and BEAM. Moreover, the amount of activation was higher in TRfail than TRsucc for most muscles (Table 1). A decrease in muscle activity during learning has previously been observed [34, 89]. Keller et al. [90] found reduced H-reflexes after a slackline training, which could explain less muscle activity with higher proficiency, due to less coactivation of agonist and antagonist muscles among a joint. In addition to this feedback-theory, we introduce a feedforward-approach. Our assumption is that synergies that are relevant for specific subtasks at a given time need to dominate over other synergies that may be activated at similar timings but are irrelevant to those subtasks. As proficiency increases and there is higher distinctness among activation coefficients, synergies for the relevant subtasks can be less activated.

The current study revealed a decrease of trial-to-trial variability during learning, and with higher proficiency (Figure 3). These findings strengthen previous studies on trial-to-trial variability as outlined in the introduction [19–22, 24, 26, 27]. Regarding the overall trial-to-trial similarity of synergy vectors, we found a transfer effect of a balancing training on the TIGHTROPE to the BEAM. However, there were no differences between startLINE and endLINE suggesting that differences did not occur due to movement-artefacts or sensor-noise. Through cluster analysis we were able to detect whether changes in variability happen in all synergy vectors and interestingly, cluster 6 did not reveal any changes in variability due to proficiency or learning. Surprisingly, in cluster 3, lag% was lowest in TIGHTROPE. An explanation could be, that in order to perform a step, regardless of the task and proficiency, activation patterns of these synergy vectors have to be quite specific and do not allow much trial-to-trial variability. On basis of our analyses, we can only speculate about this feature. The other clusters showed that trial-to-trial similarity increases with movement proficiency. While we observed an increase of trial-to-trial similarity from startBEAM to endBEAM and from TRfail to TRsucc in certain clusters, other clusters showed no changes throughout the learning process. This suggests that early learning is driven by an increase in the consistency of certain synergies, while other synergies increase their consistency during a later learning stage, i.e.: with higher proficiency levels. A noteworthy finding from the cluster analysis was that high trial-to-trial variability did not necessarily correspond to the contributions of synergies to the task. While most synergies contributed more in TIGHTROPE, cluster 4 - primarily formed by shank muscles - actually contributed more in LINE. Interestingly, despite its higher contribution in LINE, cluster 4 also exhibited the highest trial-to-trial variability in TIGHTROPE (Figure 5).

**Figure 5:**
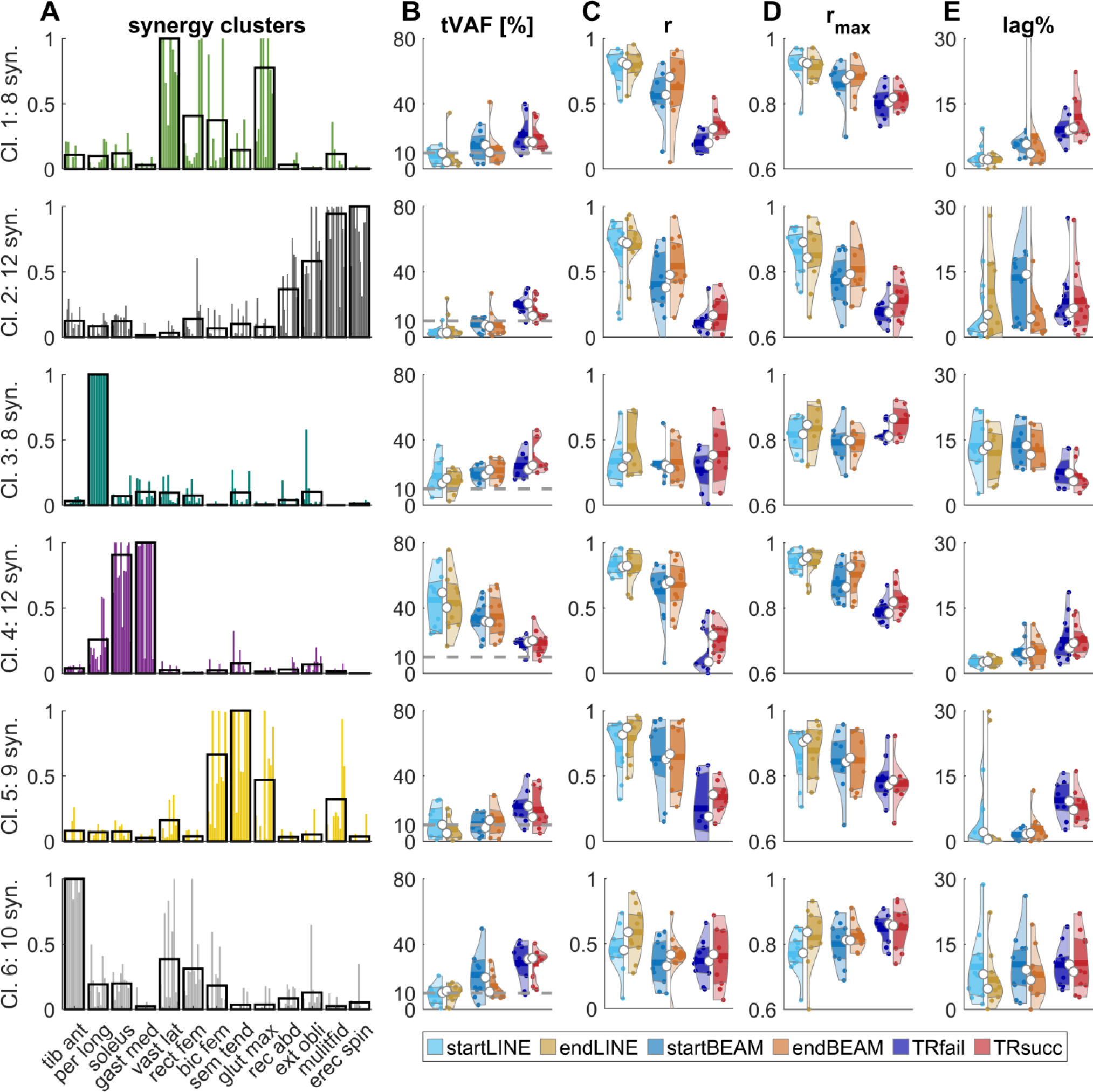
**A:** muscle weightings of clustered synergies. Black borders are the cluster (Cl.) centroids, and colored bars (similar to Figure 4) represent the synergy vectors (syn.) that belong to this cluster. **B-E:** Violin plots represent the total variance accounted for (tVAF), pearson correlation coefficient (r), cross-correlation coefficient (r_max_) and the lag-time (lag%) for each cluster. Violin plots: each colored circle represents one participant; thick lines represent mean values; white circles indicate median values; dark areas indicate quartiles.

For a more comprehensive understanding of changes in trial-to-trial variability, we also examined variability of EMG envelopes and joint angles (Figure 3). Overall EMG and joint angle variability were similar to overall synergy variability regarding task proficiency. Surprisingly, overall trial-to-trial similarity of kinematic data was not only higher with proficiency and after learning, but also in endLINE compared to startLINE. Therefore, we hypothesize that synergies reflect motor planning through the central nervous system, while kinematics are more affected by peripherical noise in the movement execution [18, 19].

Stance phases duration differed between tasks and between TRsucc and TRfail. Namely, stance phases were shorter in TRsucc than TRfail (supplementary material). This explains the smoother synergy activation patterns in LINE and BEAM compared to TIGHTROPE [32] (Figure 6). One could assume higher trial-to-trail variability in TIGHTROPE as a result of less smoothed activation coefficients, but this would not explain variability differences between LINE and BEAM, as stance duration was not different between these two tasks. To further evaluate if our findings were affected by the different task durations, we modified the low-pass cutoff frequency for each trial, based on its duration and repeated our main analyses on synergies. Detailed information and results for the additional analyses are provided in supplementary materials. Briefly, these analyses showed similar results according to trial-to-trial similarity and distinctness between the tasks. However, when comparing TRfail to TRsucc, not only trial-to-trial similarity, but also distinctness of activation coefficients revealed an increase. In summary, we drew the same conclusions based on the additional and the main analyses. Namely, fine tuning of synergy recruitment, i.e. increasing trial-to-trial similarity and activation distinctness, is important for motor learning. We hypothesize that after a more completed learning process (i.e. all attempts of TIGHTROPE walking are successful) both will increase even more and precede similar levels like BEAM and LINE.

**Figure 6:**
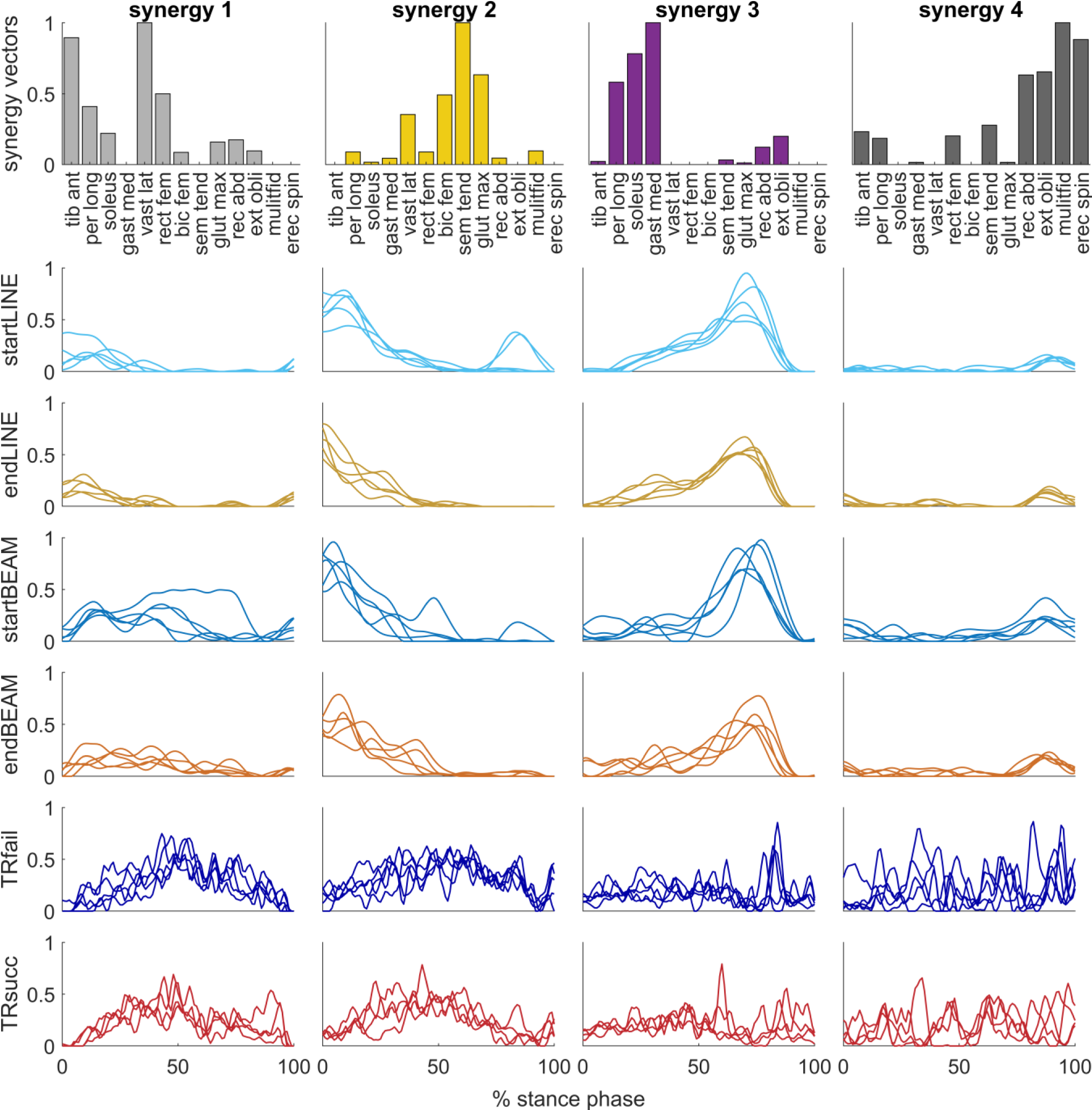
All extracted synergy vectors (bar plots) and corresponding activation coefficients (waveform plots in the same column) for each condition of one participant (P8). Each waveform represents the activation coefficient of one trial. Bar colors indicate the cluster, which the motor module belongs to, and are the same as in Figure 4 and Figure 5.

**Figure 7:**
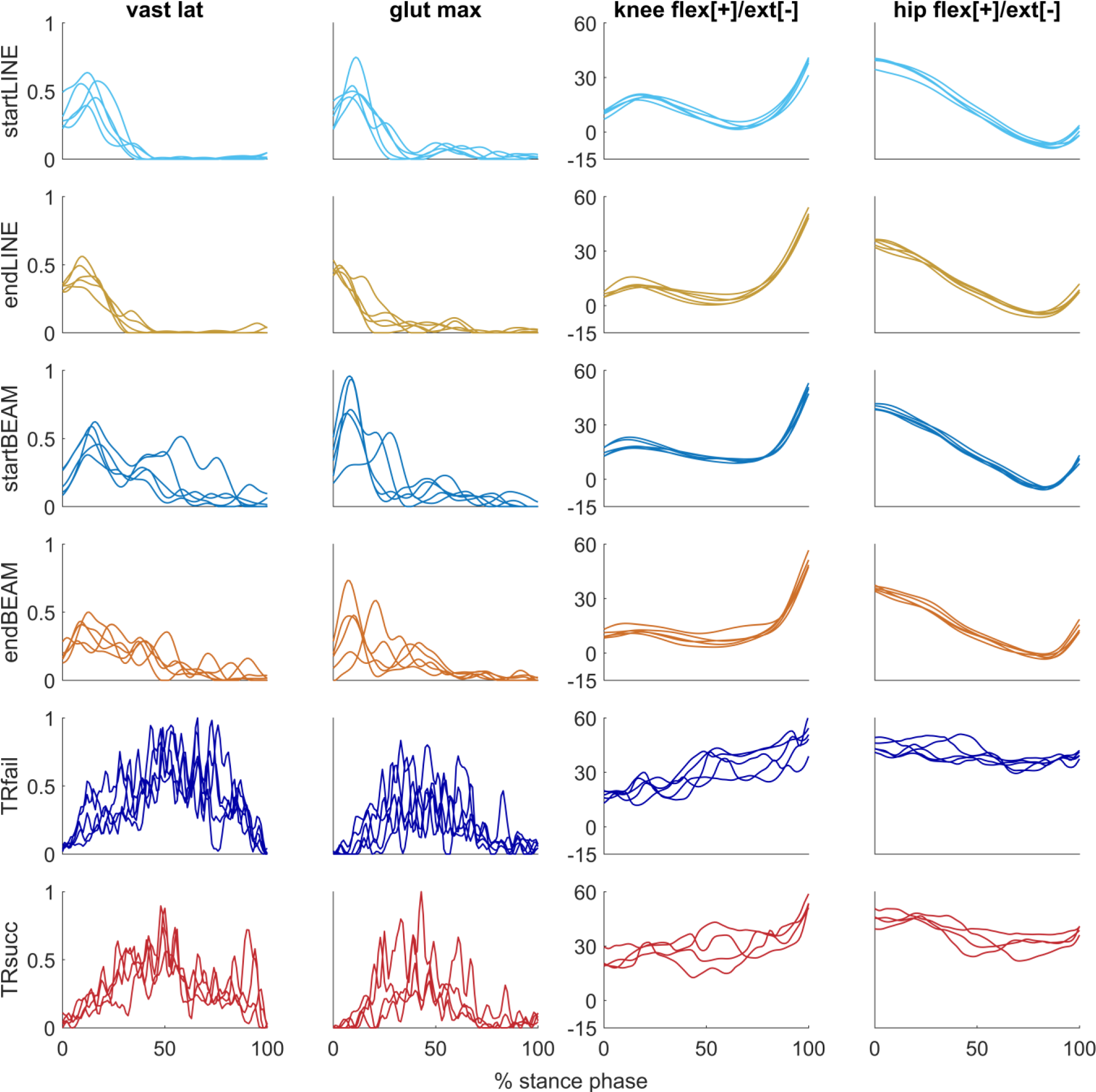
Muscle activation (example of two muscles) and joint angle waveforms (example of two joint angles) from one participant (P8). Each waveform represents one trial per condition. vast lat = vastus lateralis; glut max = gluteus maximus; flex = flexion; ext = extension.

In the field of motor learning and development three theories are widely discussed [24]. The strict nativist view proposes that locomotor modules remain robustly conserved into adulthood, supported by the spatial synergy model [36, 52] and studies observing basic stepping patterns in newborns [91]. The learning hypothesis suggests that unstructured movement patterns are transformed into structured solutions during development through the interaction between the body and the environment, evidenced by studies showing high trial-to-trial variability in early learning [19–22, 24, 26, 27]. A combined approach posits the existence of conserved movement patterns enriched with new patterns to represent a wider range of tasks. This concept has been recently supported by muscle synergy analysis in locomotion development [24]. In line with this, Cheung et al. [92] observed both, consistent and variable synergies during running development. Here, we found similar synergy vectors across tasks. In a subsequently analysis we confirmed this finding, by extracting synergy vectors separately for each condition. Briefly we found that similar motor control was utilized for all tasks. A more detailed discussion of this analysis is provided in the supplementary material. Beside similar synergy vectors, we observed higher variability in their activations in low proficiency levels. Furthermore, certain synergy vectors showed minimal contribution to LINE and BEAM tasks but were important for TIGHTROPE, indicating an enrichment of the motor control repertoire. These findings provide support for the combined nativist and learning theory.

Our study included two notable limitations. Firstly, due to the intra-session design, we captured a limited number of gait cycles per condition. Oliveira et al. [35] suggested to extract muscle synergies over a minimum of 20 concatenated steps to account for trial-to-trial variability in movement execution. To address this, we performed our main analysis on concatenated data of all conditions, providing a larger sample size of 24 to 30 stance phases per participant. Secondly, we considered the learning process to be complete when participants successfully walked across the entire tightrope in four out of five consecutive attempts, which may not reflect a high level of proficiency. Nonetheless, despite this limitation, we observed significant changes from TRfail to TRsucc in most analyzed parameters.

In summary, our study aimed to investigate motor learning using synergy analysis through a within-session, within-participant study design. We found that increasing movement proficiency led to higher distinctness and trial-to-trial similarity of synergy activation coefficients. Our findings suggest that freezing the number of degrees of freedom in early learning is a result of higher temporal overlap of synergy recruitment. Furthermore, our results support the notion that variability during the learning process increases the likelihood of finding the optimal motor command. We conclude that finetuning of synergy recruitment is crucial for motor learning.

## 6 Acknowledgements

We thank Francois Hug for his fruitful discussion and feedback on our synergy and data analysis.

## 7 Author contributions

P. K.: Conceived and designed research, performed experiments, Analyzed data, Interpreted results of experiments, Prepared figures, Drafted manuscript, Approved final version of manuscript

W. K.: Conceived and designed research, Edited and revised manuscript, Approved final version of manuscript

B. G.: performed experiments, Interpreted results of experiments, Approved final version of manuscript

E. W.: Conceived and designed research, Performed experiments, Approved final version of manuscript

A. B.: Conceived and designed research, Approved final version of manuscript

H. K.: Conceived and designed research, Performed experiments, Interpreted results of experiments, Edited and revised manuscript, Approved final version of manuscript

