## supplementary material for "Trial-to-trial similarity and distinctness of muscle synergy activation coefficients increases during learning and with a higher level of movement proficiency"

#### 1 Synergy extraction – spatial synergy model

As mentioned in the main paper, the spatial synergy model describes EMG signals as a linear combination of fixed synergy vectors W and time dependent activation coefficients C [1-3]. The mathematical concept behind this, is presented in equation (1), where E is the EMG matrix, C the activation coefficient, W the synergy vector and e the residual error. Subscript *mus* indicates the number of muscles (here 13) and *tps* the number of timepoints (here the number of all trials per participant multiplied with 101). Note that *k* represents the number of extracted synergies and ranges from 1 to 12 (mus -1) in the current study, while *g* represents the synergy number (1 to *k*). The NNMF algorithm [4-6] aims to obtain the smallest possible residual error in this equation by updating C and W over numerous iterations.

$$(1) \quad E_{mus \times tps} = \sum_{g=1}^{k} C(g)_{k \times tps}\, W(g)_{mus \times k} + e$$

We used an advanced non-negative-matrix-factorization (NNMF) algorithm introduced by Kim & Park [7] based on the block principal pivoting method for the non-negativity constrained least squares problem. In this method, convergence is reached as a stop criterium, in contrast to the classic NNMF method which could get stuck in local minima. Hence, we used the "nmf_bpas" octave function where 50 to 5000 iterations were allowed, to reach a convergence criterium of $10^{-5}$ for f in equation (2). Both, α and β are the mean of E (but one could give each an individual initial guess instead), and the subscript F indicates the Frobenius norm.

$$(2) \quad f(W,C) = \frac{1}{2}(\|E - WC\|_F^2 + \alpha\, \|W\|_F^2 + \beta \|C\|_F^2$$

It has been shown, that the number of needed iterations at which a NNMF algorithm converges and its final solution is strongly affected by the initialization (= first guess) of W and C, that are - if not specified - random inputs [8-10]. Recently, initialization methods like single-value-decompensation (SVD) principal-component-analysis, or spatial distributions gained some attention in muscle synergy analyzes due to their positive effects of hastening the NNMF algorithm and performing better when activation coefficients of the different synergies are more correlated [2, 11]. Therefore, we here used the NNSVDLRC (nonnegative single-value-decompensation with low-rank correction) function introduced by Atif et al. [9] with default inputs (stop criterion: 0.05; maximum number of iterations: 20) to obtain better initial guesses for W and C. This algorithm was designed for an improved performance on low ranks (k) which are important in muscle synergy analysis.

#### 2 k-means clustering

As mentioned in the main paper, we used octave's in-built "kmeans" function to cluster similar synergies among participants. The following properties were applied: squared Euclidean distance; k-means++ initialization algorithm; maximum iteration number of $10^{100}$ to reach a change of any centroid less than 0.0001; 5000 replicates.

#### 3 Task duration

##### 3.1 Methods

Previous studies found shorter gait-cycle durations after locomotor development [27-29]. To evaluate if stance-phase durations were shorter with higher proficiency and after a learning process, durations of all trials within one condition were averaged and compared across conditions with a 2-way ANOVA (see main paper statistics).

##### 3.2 Results

A significant effect of TASK ($p < 0.001$), TIME ($p < 0.05$) and the interaction TASK × TIME ($p < 0.05$) was found on the duration of stance phases (Figure 1). TIGHTROPE had longer stance phases ($p < 0.001$) than LINE and BEAM, with no difference between the latter two. Contrasts showed significantly shorter stance phases in TRsucc than TRfail ($p < 0.05$) (Figure 1).

#### 4 EMG – trial specific low pass cutoff frequency

##### 4.1 Methods

Reviews by Hug et al. [30] and Turpin et al. [2] call for greater awareness of the choice of cutoff frequencies for low-pass filters, prior to synergy extraction. Simply put, a lower cutoff frequency leads to a smoother EMG envelope. This 'wider' activation profile contains less variation and probably more overlap between different muscle profiles, which naturally affects synergy results. Several studies have investigated the effect of different low-pass filters on extracted motor modules. For instance, lower cutoff frequencies led to a higher total variance accounted for (tVAF) at a given number of synergies, which consequently also affected the choice of the required synergy number (NoS) for a movement[30-34]. However, Hug et al. [31] showed, that NoS was not affected by different cutoff frequencies, when the knee-point method was applied (as also used in the current study) in contrast to fixed thresholds (e.g. tVAF ≥ 90%). Moreover, low-pass filters also altered synergy vectors W and activation coefficients C extracted via NNMF [33, 34]. This is not only a problem when various studies are compared, but also matters when movements with different duration times are investigated within the same study [30]. To address this issue recent studies tried to gain similar smoothed electromyography (EMG) profiles by determining the low-pass cutoff frequency in relation to movement duration, e.g. 5 – 12 Hz for 60 – 140% of an optimal pedaling rate according to pedaling rates [20], 9 Hz for walking and 12 Hz for pedaling based on a machine learning pattern recognition algorithm [35], or by dividing a fixed cutoff frequency by the trial specific duration, i.e. 3.5 Hz/duration for treadmill walking [36] and 7 Hz/duration for overground walking of post-stroke patients [37]. For the current study, we used the same procedure as Banks et al. [37] and determined a trial specific cutoff frequency for the low-pass filter - 7 Hz/trial-duration. All other EMG processing and synergy extraction steps were done the same way as our main analysis (main paper). We aimed to determine whether the different durations would alter our main synergy results in terms of complexity, distinctness, and trial-to-trial variability. Figure 2 shows, how different cutoff frequencies affect EMG smoothing.

##### 4.2 Results

An average of 5.6 ± 2.22 NoS was determined among participants. For tVAF1 a significant effect of TASK ($p < 0.001$) was observed, with highest tVAF1 in TIGHTROPE, followed by BEAM and LINE at last ($p < 0.001$). There was also a significant effect of TASK in tVAFNoS ($p < 0.05$) which was higher in TIGHTROPE compared to LINE ($p < 0.05$). Regarding the distinctness of activation coefficients, the ANOVA revealed a significant effect of TASK for r ($p < 0.02$), $r_{max}$ and %lag (p <

0.001). Activation coefficients were more correlated to each other (r: $p < 0.01$; $r_{max}$: $p < 0.001$) in TIGHTROPE compared to LINE and BEAM. The lag% was higher in LINE than BEAM ($p < 0.05$) and lowest in TIGHTROPE ($p < 0.001$). Additionally, $r_{max}$ was significantly affected by the interaction TASK × TIME ($p < 0.05$), where contrasts revealed a decrease during learning on the TIGHTROPE ($p < 0.05$) (Figure 3).

Regarding trial-to-trial similarity, there was a significant effect of TASK ($p < 0.001$) on r, with highest correlations in LINE, followed by BEAM ($p < 0.01$) and TIGHTROPE at last ($p < 0.001$). Correlation was also significantly affected by TIME ($p < 0.05$), where contrasts revealed an increase during learning on the TIGHTROPE ($p < 0.05$). There were no significant differences in cross-correlations $r_{max}$ and %lag (Figure 3, Figure 4).

#### 5 Muscle synergy extraction from each condition independently

##### 5.1 Methods

To investigate whether similar motor modules were utilized across different walking conditions, we performed individual synergy extraction for each condition using the procedures described in the main paper. For this purpose, we concatenated the EMG matrices within each condition, and employed NNSVDLRC and NNMF. We performed two widely used analyses on this data. In the first step of our analysis, we compared synergy vectors (W) across all possible pairs among conditions using the Pearson correlation coefficient (r). A pair of synergy vectors W was considered as similar (= shared), if $r > 0.684$, which corresponds to the critical value of r for 13 muscles at $p = 0.01$ [12-18]. To account for variations in the number of synergies (NoS) across conditions and participants, we visualized the number of shared synergies ($n_{shared}$) as percentage. The percentage of shared synergies ($\%n_{shared}$) was calculated using equation (3), where subscripts *condition1* and *condition2* indicate the two compared conditions (e.g., startLINE and endLINE).

$$(3) \quad \%n_{shared} = 100 \, \frac{n_{shared}}{min\,(NoS_{condition1},\ NoS_{condition2})}$$

In the second step, we reconstructed synergy activation coefficients (C) for all conditions using the synergy vectors (W) from either startLINE or startBEAM. We employed a commonly used reconstruction algorithm [17, 19-24] based on the updating rule for NNMF proposed by Lee and Seung [5], as described in equations (4). W from one condition was held fixed (suffix: fix) to reconstruct C (suffix: rec), with the corresponding EMG matrix (E) from another condition. After an initial random guess for the reconstruction matrix, numerous iterations (n, here ranging from 50 to 5000) were made until the function f(W,C) reached a convergence criterion ($10^{-5}$) described in equation (5). Subscripts i and j indicate the row and column, while superscript T indicates the transposed matrix. Crec was used to calculate the reconstructed tVAF (tVAFrec), together with E and Wfix. Consistent with previous studies [24-26], synergy vectors were assumed to be similar across conditions if the tVAFrec exceeded 80%.

$$(4) \quad Crec_{ij}^{(n)} = Crec_{ij}^{(n-1)} \left( \frac{(Wfix^T\,E)_{ij}}{(Wfix^T\,Wfix\,Crec^{(n-1)})_{ij}} \right);$$

$$(5) \quad f(Wfix, Crec) = \frac{\|E - Wfix\,Crec\|_F}{\sqrt{m\,n}}$$

#### 5.2 Results

**Figure 5 shows the high amount of shared synergy vectors across the LINE and BEAM tasks for all participants. A high range, and on average a smaller %$n_{shared}$ was found between LINE or BEAM conditions with TIGHTROPE conditions across participants. Reconstruction procedures revealed, that tVAFrec was > 80% in all participants, for all LINE and BEAM conditions. In contrast, only six participants exceeded the threshold in TIGHTROPE conditions, if activation coefficients were reconstructed by synergy vectors of startLINE. If activation coefficients were reconstructed by startBEAM, tVAFrec of one participant was under our criterium for TRfail and two participants for TRsucc (**

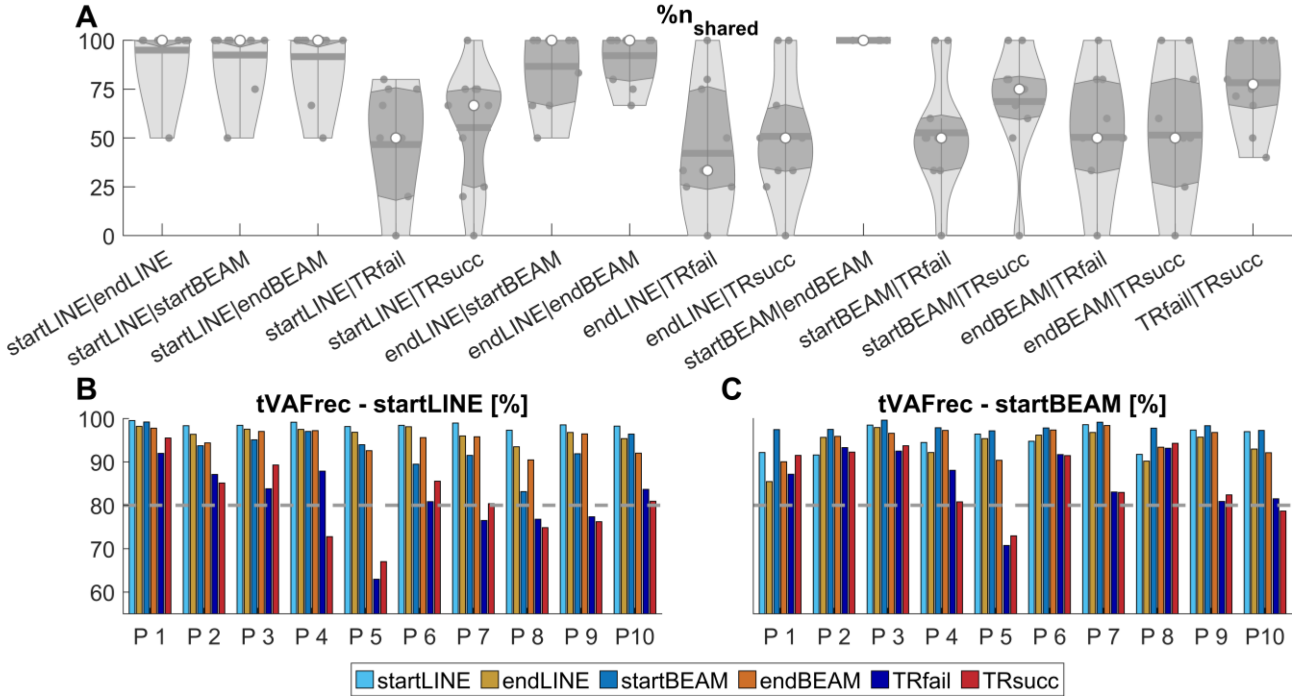


Figure 5).

#### 5.3 Discussion

Here we want to deeper discuss our approach on extracting synergies over different conditions. Similar tVAFNoS values (main paper) indicate, that extracted synergies reflect EMG variability equally among conditions. However, computing muscle synergies over different conditions has previously been done only in few studies [12, 20, 38-40]. We here utilized this approach due to the following four considerations: (1) similar movement goals are controlled via similar muscle synergies, (2) the small number of trials within each condition, (3) computing trial-to-trial similarity among similar synergy vectors, and (4) the computational problem of extracting accurate synergies, if activation coefficients timing overlaps.

(1) As outlined in the introduction, an important feature of motor control is the recruitment of similar synergy vectors for similar mechanical goals across various tasks [16, 18, 20, 25, 41, 42]. In the current study, the same movement goal – performing a step – was intended in all conditions. Therefore, we hypothesized that similar synergy vectors were used. In an additional analysis, synergies were extracted from each condition independently [20, 39, 40] to verify this assumption. In a first step, we analyzed the percentage of shared synergy vectors (r > 0.684 [12-18]). While a high

percentage of vectors was shared among LINE and BEAM conditions, less synergies were shared if LINE and BEAM conditions were compared to TRfail or TRsucc (Figure 5). In a second step, we reconstructed the activation coefficients of all conditions with synergy vectors of startLINE or startBEAM. We found sufficient reconstruction performance (tVAF > 80% [24-26]) for all LINE and BEAM conditions (Figure 5). When reconstructing TIGHTROPE conditions from startLINE, 6 participants achieved a tVAF above 80%, while for startBEAM, it was achieved by 9 participants in TRfail and 8 participants in TRsucc. Turpin et al. [43] found that some synergies were barely activated in cycling with low exercise intensity, which could lead to poor performances of factorization methods in detecting them [2, 44]. Similarly, in our study, we observed differences in the contribution of synergy vectors across conditions (see main paper). Cluster 2, primarily composed of trunk muscles, had a tVAF below 10% for LINE and BEAM conditions but exceeded the threshold for TIGHTROPE. This may explain the relatively lower reconstruction accuracy of TIGHTROPE compared to LINE and BEAM conditions since these synergies were probably not captured by extracting synergies separately. Furthermore, cluster 1, predominantly composed of quadriceps and gluteus muscles, exhibited a tVAF below 10% in most LINE synergies but only in some BEAM synergies, which could explain higher reconstruction accuracies of TIGHTROPE when using startBEAM rather than startLINE. The reconstruction results, combined with cluster analyses, indicate that there is a presence of similar synergy vectors across tasks, but additional synergies are either added or more activated during balancing tasks. The low percentage of shared synergies in some participants may be due to the difficulty of accurately extracting synergy vectors when their activation timing overlaps (see below).

(2) Several studies proposed the importance of variability in EMG data by concatenated trials for synergy computation [2, 45-47]. Oliveira et al. [45] suggested to compute muscle synergies over a minimum of 20 concatenated gait-cycles. Due to the within-session design of the current study, each condition only included four to five stance-phases. Concatenating data of all conditions resulted in a total of 24 to 30 stance phases per participants.

(3) To ensure that the variability of activation coefficients could be meaningful quantified, the same synergy vectors were used across conditions. This is in line with Cheung et al. [48] who first clustered synergy vectors among bowling sessions and reconstructed EMG matrices of each session with the cluster centroids.

(4) Increasing the time overlap (= correlation) of synergy activation coefficients in simulated or real datasets, decreases the accuracy of extracted synergy vectors. Namely, with sufficient coupling, synergy vectors merged, due to underlying assumptions of factorization algorithms [11, 44, 46]. In our opinion, the distinctness and timing of activation coefficients might reflect an essential feature for movement proficiency and learning. Calculating muscle synergies over all conditions may overcome limitations of extracting algorithms.

With this additional analysis we showed that similar synergies were utilized through the tasks. Moreover, we hypothesize that calculating synergies over different tasks with similar movement goals (i.e.: performing a step), provides salient information in synergy analysis.

### 6 Results of individual muscle and joints

#### 6.1 Muscle and joint abbreviations

Muscles: tibialis anterior (tib_abt), peroneus longus (per_long), soleus, gastrocnemius medialis (gast_med), vastus lateralis (vast_lat), rectus femoris (rect_fem), biceps femoris (bic_fem), semitendinosus (sem_tend), gluteus maximus (glut_max), rectus abdominis (rect_abd), extensor obliques (ext_obli), multifidus (multifid) and erector spinae iliocostalis (erec_spin).

Joints: ankle plantar-/dorsiflexion (ankle_flex), knee flexion/extension (knee_flex), hip flexion/extension (hip_flex), hip ab-/adduction (hip_ad), hip internal/external rotation (hip_rot), lumbar flexion/extension (lumb_flex), lumbar medial/lateral bending (lumb_bend), and lumbar internal/external rotation (lumb_rot).

#### 6.2 Muscle activations individual

Trial-to-trial similarity in muscle activation patterns measured by Pearson correlation coefficient (r) was significantly affected by TASK in all muscles apart from rec_abd and ext_obli (tib_ant, per_long: $p < 0.05$; rect_fem: $p < 0.01$; others: $p < 0.001$). Trial-to-trial similarity was smaller for TIGHTROPE than BEAM in eight muscles (vast_lat: $p < 0.05$; soleus, gast_med, bic_fem, sem_tend, glut_max, multifid, erec_spin: $p < 0.001$), and smaller for TIGHTROPE compared with LINE in all muscles (tib_ant, per_long: $p < 0.05$; rect_fem: $p < 0.01$; others: $p < 0.001$). Trial-to-trial similarity was smaller for BEAM compared to LINE in five muscles (soleus, rect_fem: $p < 0.05$; multifid: $p < 0.01$; gast_med, erec_spin: $p < 0.001$). Additionally, similarity was significantly affected by TIME (per_long, gast_med, glut_max: $p < 0.05$), and TASK × TIME (gast_med, glut_max: $p < 0.05$; soleus: $p < 0.01$) in three muscles. Contrasts revealed higher similarities in endBEAM than startBEAM (gast_med, multifid: $p < 0.05$, erec_spin: $p < 0.01$) and TRsucc than TRfail (per_long, gast_med, glut_max: $p < 0.05$) (Table 1).

Trial-to-trial similarity measured by cross-correlation coefficient ($r_{max}$) was significantly affected by TASK in six muscles (soleus, gast_med, glut_max, ext_obli, multifid, erec_spin: $p < 0.001$), where TIGHTROPE similarity was smaller than BEAM (soleus: $p < 0.05$; others: $p < 0.001$) and LINE ($p < 0.001$). BEAM similarity was smaller than LINE in four muscles (soleus: $p < 0.05$; gast_med, multifid, erec_spin: $p < 0.001$). Additionally, similarity was also significantly affected by TIME in four muscles (per_long, soleus, gast_med, multifid, erec_spin: $p < 0.05$) with higher similarities in END, and TASK × TIME in soleus ($p < 0.05$). Contrasts revealed higher similarities in endLINE than startLINE (erec_spin: $p < 0.05$), endBEAM than startBEAM (soleus, gast_med, erec_spin: $p < 0.05$) and TRsucc than TRfail (multifid: $p < 0.05$; soleus: $p < 0.01$) (Table 2).

The lag% was significantly affected by TASK for all muscles apart from multifid and erec_spin (tib_ant, per_long, soleus, rec_abd: $p < 0.05$: gast_med; rect_fem, bic_fem, multifid: $p < 0.01$; others: $p < 0.001$). lag% was significantly lower in TIGHTROPE than BEAM for per_long (per_long: $p < 0.05$). For all other muscles TIGHTROPE had a higher lag% than LINE (tib_ant, gast_med, rec_abd: $p < 0.05$; soleus, rect_fem, bic_fem, multifid: $p < 0.01$; others: $p < 0.001$) and for sem_tend and glut_max also than BEAM ($p < 0.001$). BEAM had higher lag% than LINE in three muscles (gast_med, bic_fem: $p < 0.05$; vast_lat: $p < 0.01$) (Table 3).

#### 6.3 Joint angles individual

Trial-to-trial similarity measured by Pearson correlation coefficient (r) was significantly affected in all joints by TASK (ankle_flex: $p < 0.01$; others: $p < 0.001$). TIGHTROPE similarity was always lower than BEAM (ankle_flex; $p < 0.05$; others: $p < 0.001$) and LINE (ankle_flex: $p < 0.01$; others: $p < 0.001$). BEAM similarity was lower than LINE in three joints (knee_flex, hip_ad: $p < 0.05$; lumb_bend: $p < 0.001$). It was also significantly affected by TIME in some joints (hip_ad, hip_rot, lumb_rot: $p < 0.05$; ankle_flex, hip_flex: $p < 0.01$), with lower similarity in START than END. Additionally, a significant effect of TIME × TASK was found in two joints (hip_rot: $p < 0.01$;

hip_flex: $p < 0.001$). Contrasts showed that r was lower in startLINE than endLINE (hip_ad: $p < 0.05$) and TRfail than TRsucc (ankle_flex: $p < 0.05$) (Table 4).

Trial-to-trial similarity measured by cross-correlation coefficient ($r_{max}$) was significantly affected by TASK in all joints (lumb_flex: $p < 0.05$; hip_flex, hip_rot: $p < 0.05$; others: $p < 0.001$). TIGHTROPE similarity was lower than BEAM in six joints (ankle_flex, lumb_flex, lumb_bend: $p < 0.05$; knee_flex, hip_ad, lumb_rot: $p < 0.001$) and LINE in all joints (hip_flex, hip_rot: $p < 0.05$; lumb_flex: $p < 0.01$; othert: $p < 0.001$). BEAM similarity was lower than LINE in lumb_bend ($p < 0.01$). It was also significantly affected by TIME in two joints (lumb_rot: $p < 0.01$; lumb_flex: $p < 0.001$), with lower similarity in START than END. Additionally, a significant effect of TIME × TASK was found in lumb_rot ($p < 0.05$). Contrasts showed that $r_{max}$ was lower in startBEAM than endBEAM (lumb_flex, lumb_bend: $p < 0.05$) and TRfail than TRsucc (lumb_flex: $p < 0.01$) (Table 5).

The lag% was significantly affected by TASK in five joints (hip_rot, lumb_flex: $p < 0.01$; hip_ad, lumb_bend, lumb_rot: $p < 0.001$). TIGHTROPE had a higher lag% than BEAM in four joints (hip_ad, hip_rot, lumb_rot: $p < 0.01$; lumb_bend: $p < 0.001$) and LINE in all five joints (hip_rot, lumb_rot: $p < 0.01$; other: $p < 0.001$). BEAM had a higher lag% than LINE in two jonts (hip_ad: $p < 0.05$; lumb_bend: $p < 0.001$). There was a significant effect of TIME in five joints (hip_ad, lumb_flex, lumb_rot: $p < 0.01$; knee_flex, hip_rot: $p < 0.001$) with higher lag% in START. Additionally, there was a significant effect of TASK × TIME in three joints (knee_flex, hip_ad: $p < 0.01$; hip_rot: $p < 0.001$). Contrasts revealed higher lag% in startLINE than endLINE in hip_ad ($p < 0.05$). For lumb_flex, startBEAM and TRfail had significantly higher lag% compared to endBEAM and TRsucc ($p < 0.01$), respectively (Table 6).

### 8 Figures

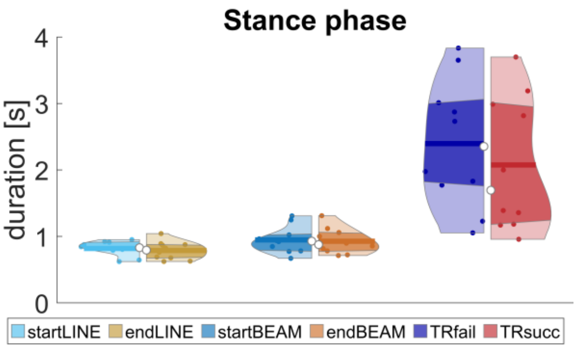


**Figure 1:** Stance phase duration (in seconds [s]) of each condition. Violin plots: each colored circle represents one participant; thick lines represent mean values; white circles indicate median values; dark areas indicate quartiles.

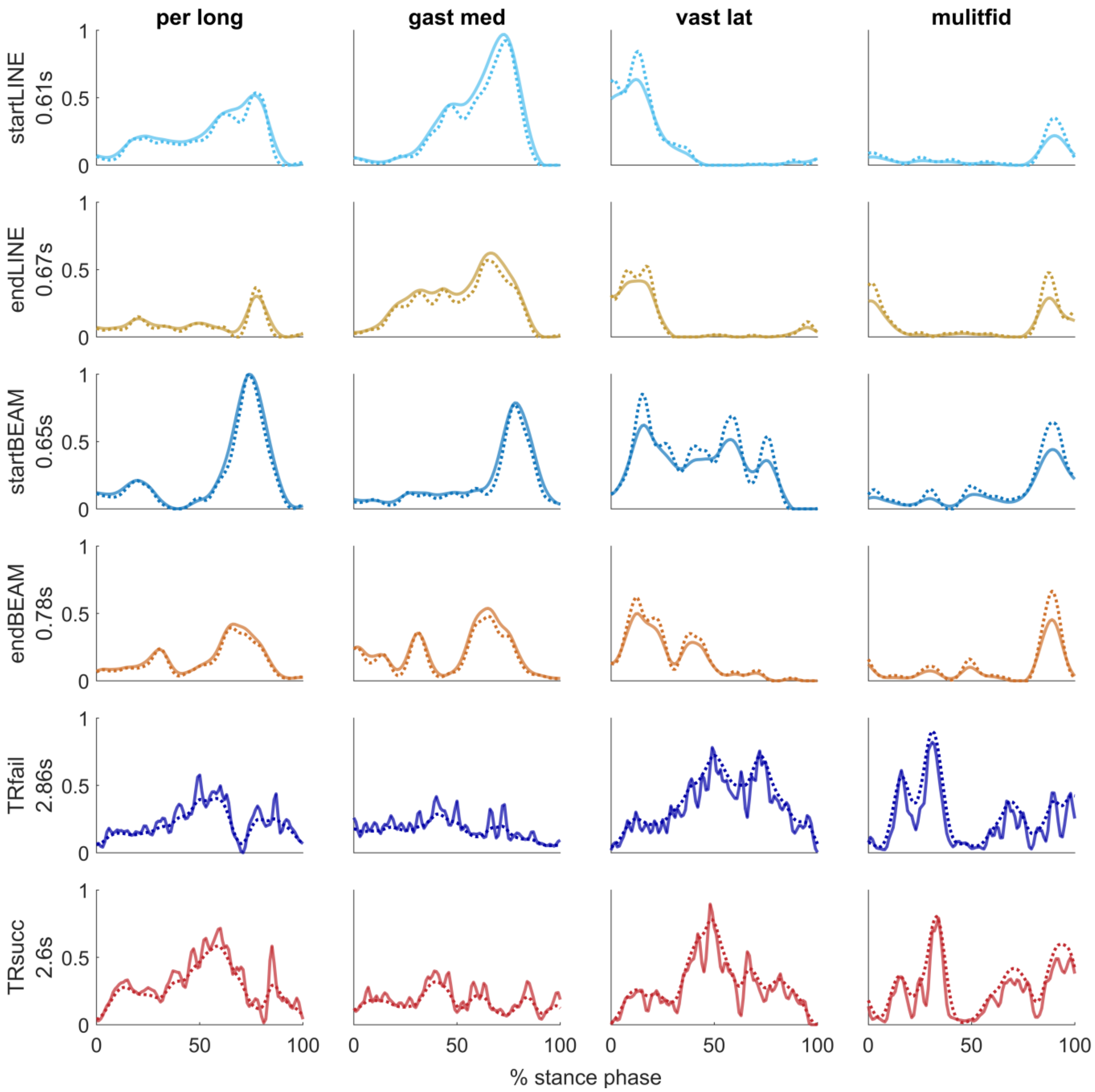


**Figure 2:** Example of the influence of different cutoff frequencies on EMG smoothing of four muscles (one trial per condition, of one participant). The solid lines represent the fixed low-pass cutoff frequency (7 Hz), and the dashed lines represent the duration-dependent cutoff frequency (7 Hz/trial duration). The duration of the trials is shown on the y axis and given in seconds [s].

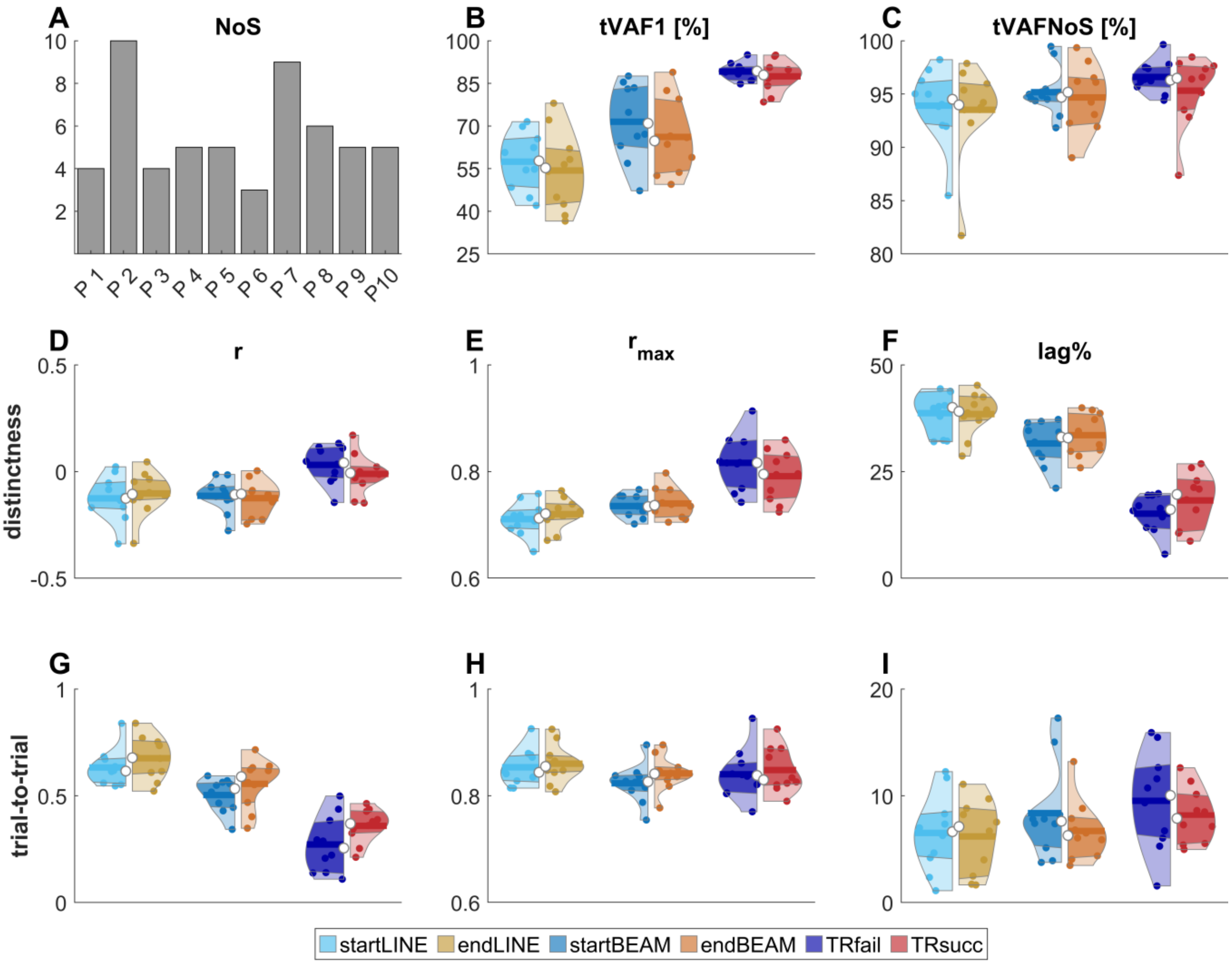


**Figure 3:** Synergies were extracted of EMG signals filtered with the duration-dependent cutoff frequencies. **A**: bars show the number of required synergies (NoS) for each participant (P1 – P10). **B-C**: the total variance accounted for one synergy (**B**: tVAF1) and NoS (**C**: tVAFNoS). **D-F**: Synergy activation coefficient distinctness and **G-I**: trial-to-trial similarity measured by Pearson correlation (**D, G**: r), maximum cross-correlation coefficient (**E, H**: $r_{max}$) and lag at $r_{max}$ (**F, I**: lag%). Violin plots: each colored circle represents one participant; thick lines represent mean values; white circles indicate median values; dark areas indicate quartiles.

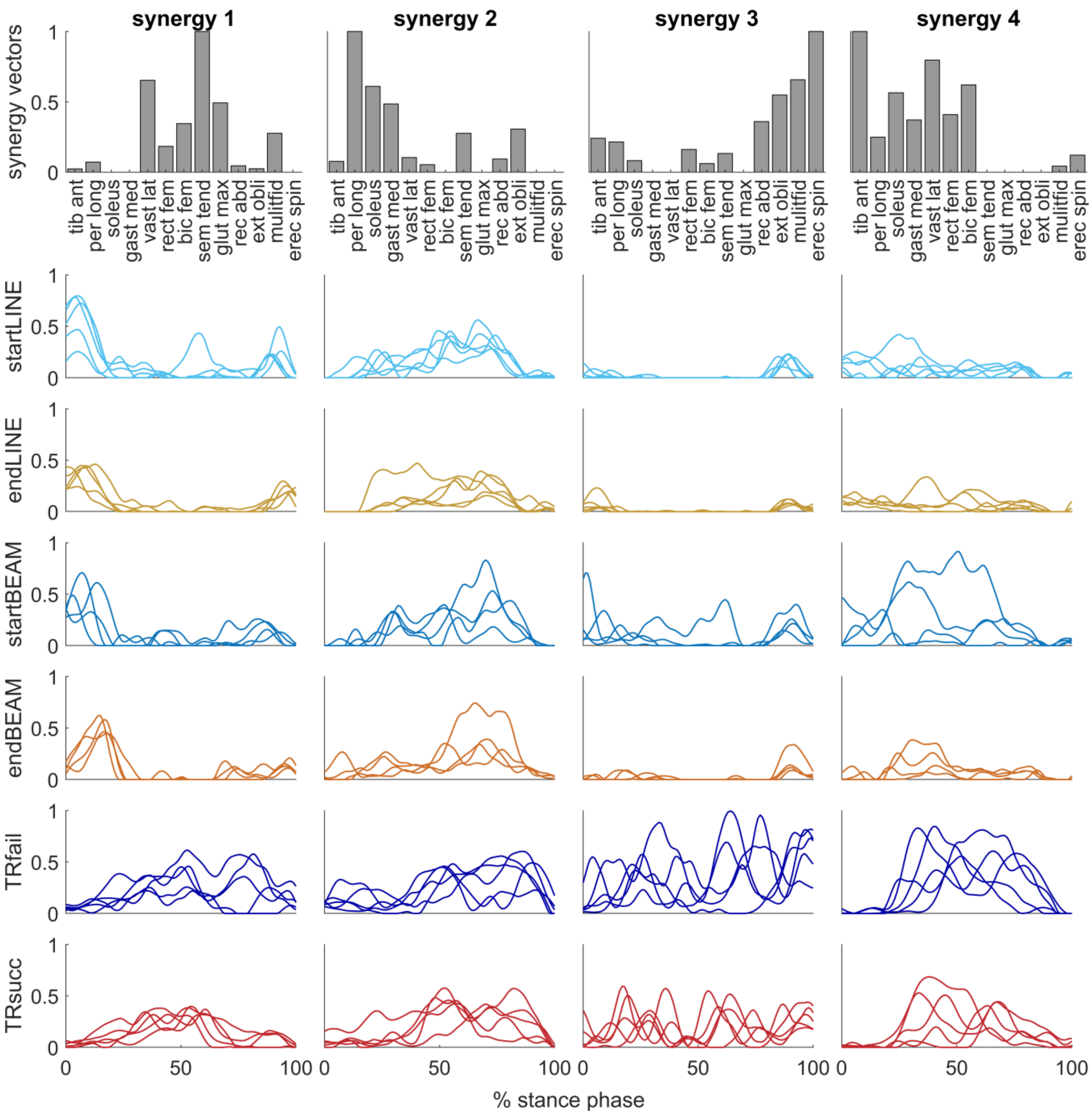


**Figure 4:** All extracted synergy vectors (bar plots) and corresponding activation coefficients (waveform plots in the same column) for each condition of one participant (P3). Each waveform represents the activation coefficient of one trial.

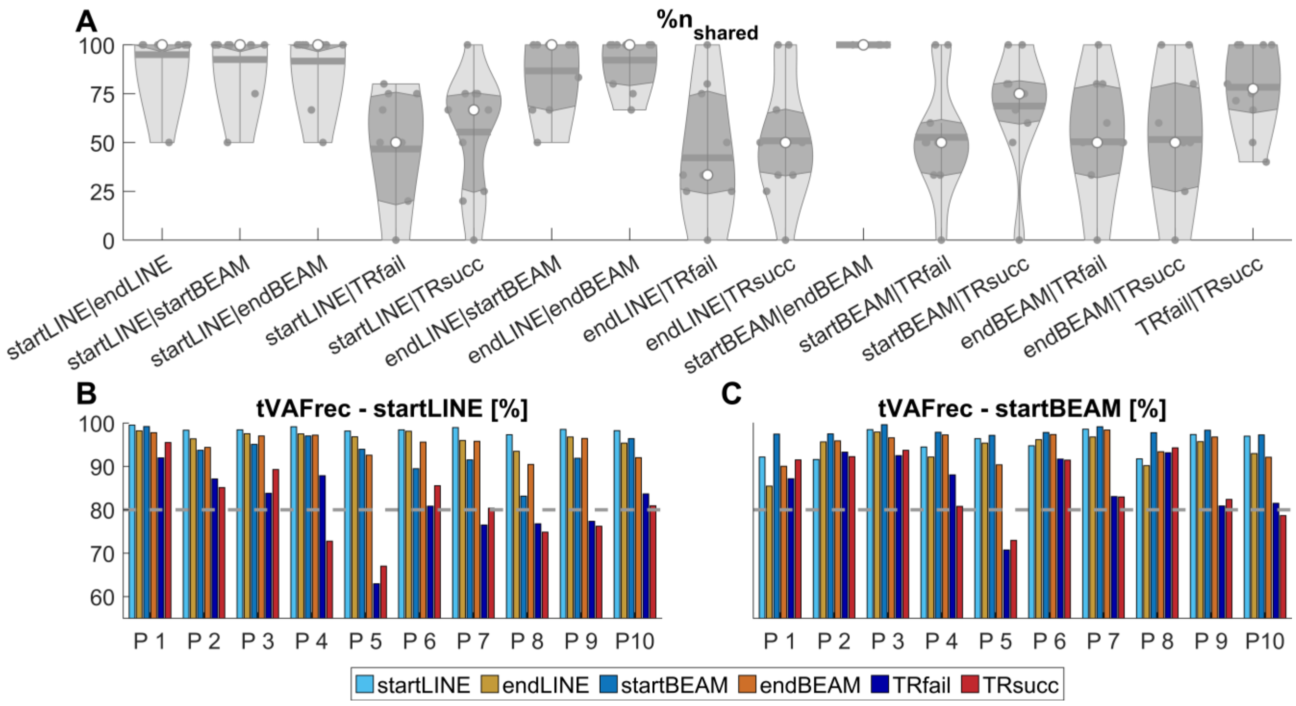


**Figure 5: A**: The percentage of shared synergy vectors (n%$_{shared}$) for all possible pairs of condition comparisons. **B-C**: the tVAF of reconstructed activation coefficients (tVAFrec) of the synergy vectors of either startLINE (**B**) or startBEAM (**C**) for all participants (P) and conditions. Violin plots: each grey circle represents one participant; thick lines represent mean values; white circles indicate median values; dark areas indicate quartiles.

### 9 Tables

**Table 1:** Mean (M) and standard deviation (SD) among participants of for trial-to-trial similarity measured by Pearson correlation coefficient r for all conditions and muscles. ANOVA revealed significant effects of TASK in all muscles apart from rec_abd and ext_obli. Significant differences observed by contrasts are indicated by *.

| | LINE | | | | BEAM | | | | TIGHTROPE | | | |
|---|---|---|---|---|---|---|---|---|---|---|---|---|
| | start | | end | | start | | end | | fail | | succ | |
| | *M* | *SD* | *M* | *SD* | *M* | *SD* | *M* | *SD* | *M* | *SD* | *M* | *SD* |
| tib_ant | 0.49 | 0.22 | 0.60 | 0.17 | 0.43 | 0.15 | 0.45 | 0.20 | 0.36 | 0.19 | 0.36 | 0.21 |
| per_long | 0.49 | 0.22 | 0.61 | 0.20 | 0.45 | 0.16 | 0.48 | 0.22 | 0.31* | 0.16 | 0.44* | 0.19 |
| soleus | 0.84 | 0.07 | 0.80 | 0.10 | 0.64 | 0.18 | 0.67 | 0.17 | 0.23 | 0.19 | 0.31 | 0.18 |
| gast_med | 0.83 | 0.08 | 0.83 | 0.08 | 0.61* | 0.21 | 0.69* | 0.19 | 0.17* | 0.16 | 0.31* | 0.11 |
| vast_lat | 0.78 | 0.20 | 0.73 | 0.28 | 0.56 | 0.23 | 0.60 | 0.20 | 0.31 | 0.18 | 0.41 | 0.16 |
| rect_fem | 0.62 | 0.23 | 0.63 | 0.31 | 0.36 | 0.28 | 0.47 | 0.33 | 0.24 | 0.13 | 0.36 | 0.18 |
| bic_fem | 0.66 | 0.20 | 0.70 | 0.18 | 0.55 | 0.29 | 0.60 | 0.16 | 0.27 | 0.17 | 0.31 | 0.11 |
| sem_tend | 0.63 | 0.27 | 0.76 | 0.17 | 0.62 | 0.24 | 0.65 | 0.18 | 0.23 | 0.14 | 0.26 | 0.16 |
| glut_max | 0.81 | 0.08 | 0.79 | 0.18 | 0.61 | 0.23 | 0.70 | 0.20 | 0.14* | 0.18 | 0.32* | 0.12 |
| rec_abd | 0.15 | 0.23 | 0.02 | 0.12 | 0.13 | 0.20 | 0.08 | 0.09 | 0.11 | 0.07 | 0.11 | 0.14 |
| ext_obli | 0.28 | 0.18 | 0.26 | 0.23 | 0.16 | 0.21 | 0.29 | 0.16 | 0.07 | 0.11 | 0.18 | 0.18 |
| multifid | 0.82 | 0.09 | 0.85 | 0.06 | 0.57* | 0.25 | 0.68* | 0.16 | 0.10 | 0.10 | 0.17 | 0.18 |
| erec_spin | 0.62 | 0.14 | 0.69 | 0.18 | 0.35* | 0.21 | 0.49* | 0.20 | 0.10 | 0.08 | 0.11 | 0.10 |

**Table 2:** Mean (M) and standard deviation (SD) among participants of for trial-to-trial similarity measured by the maximum cross-correlation coefficient $r_{max}$ for all conditions and muscles. ANOVA revealed significant effects of TASK in soleus, gast_med, glut_max, ext_obli, multifid and erec_spin. Significant differences observed by contrasts are indicated by *.

| | LINE | | | | BEAM | | | | TIGHTROPE | | | |
|---|---|---|---|---|---|---|---|---|---|---|---|---|
| | start | | end | | start | | end | | fail | | succ | |
| | *M* | *SD* | *M* | *SD* | *M* | *SD* | *M* | *SD* | *M* | *SD* | *M* | *SD* |
| tib_ant | 0.81 | 0.06 | 0.83 | 0.06 | 0.85 | 0.04 | 0.86 | 0.04 | 0.87 | 0.04 | 0.87 | 0.07 |
| per_long | 0.88 | 0.05 | 0.90 | 0.04 | 0.87 | 0.05 | 0.87 | 0.06 | 0.88 | 0.02 | 0.90 | 0.03 |
| soleus | 0.96 | 0.02 | 0.95 | 0.02 | 0.91* | 0.04 | 0.93* | 0.03 | 0.87* | 0.04 | 0.90* | 0.03 |
| gast_med | 0.95 | 0.03 | 0.95 | 0.03 | 0.88* | 0.04 | 0.92* | 0.04 | 0.84 | 0.03 | 0.86 | 0.04 |
| vast_lat | 0.91 | 0.05 | 0.89 | 0.08 | 0.85 | 0.08 | 0.87 | 0.06 | 0.86 | 0.05 | 0.86 | 0.06 |
| rect_fem | 0.85 | 0.09 | 0.85 | 0.08 | 0.80 | 0.07 | 0.84 | 0.09 | 0.82 | 0.06 | 0.84 | 0.06 |
| bic_fem | 0.83 | 0.06 | 0.84 | 0.07 | 0.84 | 0.05 | 0.83 | 0.05 | 0.82 | 0.04 | 0.80 | 0.04 |
| sem_tend | 0.86 | 0.07 | 0.89 | 0.05 | 0.85 | 0.08 | 0.86 | 0.03 | 0.83 | 0.05 | 0.80 | 0.07 |
| glut_max | 0.92 | 0.04 | 0.92 | 0.04 | 0.88 | 0.06 | 0.90 | 0.05 | 0.77 | 0.06 | 0.79 | 0.08 |
| rec_abd | 0.72 | 0.07 | 0.72 | 0.06 | 0.74 | 0.07 | 0.72 | 0.05 | 0.69 | 0.04 | 0.71 | 0.08 |
| ext_obli | 0.82 | 0.04 | 0.82 | 0.04 | 0.81 | 0.05 | 0.82 | 0.03 | 0.72 | 0.05 | 0.76 | 0.05 |
| multifid | 0.90 | 0.05 | 0.92 | 0.03 | 0.83 | 0.07 | 0.86 | 0.05 | 0.72* | 0.06 | 0.76* | 0.07 |
| erec_spin | 0.83* | 0.05 | 0.89* | 0.07 | 0.76* | 0.08 | 0.80* | 0.07 | 0.67 | 0.03 | 0.68 | 0.07 |

**Table 3:** Mean (M) and standard deviation (SD) among participants of for trial-to-trial similarity measured by the lag time lag% at the maximum cross-correlation coefficient for all conditions and muscles. ANOVA revealed significant effects of TASK in all muscles apart from multifid and erec_spin.

| | **LINE** | | | | **BEAM** | | | | **TIGHTROPE** | | | |
|---|---|---|---|---|---|---|---|---|---|---|---|---|
| | **start** | | **end** | | **start** | | **end** | | **fail** | | **succ** | |
| | *M* | *SD* | *M* | *SD* | *M* | *SD* | *M* | *SD* | *M* | *SD* | *M* | *SD* |
| tib_ant | 6.06 | 7.58 | 5.55 | 5.02 | 5.60 | 4.01 | 6.64 | 4.88 | 9.26 | 5.93 | 10.24 | 7.67 |
| per_long | 6.84 | 3.79 | 5.54 | 4.41 | 7.62 | 3.88 | 7.06 | 3.44 | 5.13 | 3.06 | 3.82 | 2.20 |
| soleus | 1.98 | 0.63 | 2.95 | 1.46 | 4.58 | 1.11 | 3.66 | 2.13 | 6.95 | 4.92 | 6.09 | 3.09 |
| gast_med | 2.94 | 1.18 | 2.72 | 0.81 | 5.63 | 2.33 | 4.39 | 1.76 | 6.68 | 4.49 | 6.56 | 3.79 |
| vast_lat | 2.89 | 3.32 | 3.39 | 7.39 | 7.09 | 7.59 | 5.04 | 3.76 | 6.93 | 2.86 | 7.99 | 4.22 |
| rect_fem | 3.30 | 3.89 | 6.44 | 11.24 | 7.06 | 7.24 | 9.08 | 11.58 | 10.93 | 3.64 | 8.76 | 4.60 |
| bic_fem | 7.43 | 9.48 | 5.74 | 10.69 | 5.51 | 6.17 | 5.98 | 5.98 | 6.60 | 3.78 | 12.33 | 5.80 |
| sem_tend | 3.92 | 5.89 | 3.10 | 5.58 | 5.23 | 8.82 | 1.63 | 1.83 | 8.37 | 5.12 | 9.17 | 5.52 |
| glut_max | 2.16 | 1.10 | 3.19 | 3.85 | 5.20 | 4.17 | 5.33 | 6.02 | 13.76 | 4.57 | 11.47 | 7.26 |
| rec_abd | 17.52 | 8.52 | 20.07 | 5.64 | 16.26 | 8.22 | 17.80 | 8.88 | 11.81 | 3.78 | 12.80 | 6.11 |
| ext_obli | 8.43 | 6.32 | 8.07 | 5.82 | 7.45 | 3.80 | 8.62 | 5.34 | 8.63 | 5.16 | 9.14 | 6.11 |
| multifid | 0.90 | 0.61 | 3.78 | 6.61 | 4.95 | 5.35 | 4.32 | 7.74 | 6.98 | 4.70 | 6.22 | 6.70 |
| erec_spin | 7.94 | 8.33 | 10.15 | 14.38 | 15.95 | 12.86 | 14.14 | 17.25 | 11.02 | 7.92 | 12.78 | 8.24 |

**Table 4:** Mean (M) and standard deviation (SD) among participants of for trial-to-trial similarity measured by Pearson correlation coefficient r for all conditions and joint angles. ANOVA revealed significant effects of TASK in all joints Significant differences observed by contrasts are indicated by *.

| | **LINE** | | | | **BEAM** | | | | **TIGHTROPE** | | | |
|---|---|---|---|---|---|---|---|---|---|---|---|---|
| | **start** | | **end** | | **start** | | **end** | | **fail** | | **succ** | |
| | *M* | *SD* | *M* | *SD* | *M* | *SD* | *M* | *SD* | *M* | *SD* | *M* | *SD* |
| ankle_flex | 0.95 | 0.04 | 0.97 | 0.02 | 0.92 | 0.07 | 0.94 | 0.06 | 0.77* | 0.21 | 0.88* | 0.06 |
| knee_flex | 0.97 | 0.02 | 0.97 | 0.02 | 0.93 | 0.07 | 0.91 | 0.07 | 0.53 | 0.31 | 0.54 | 0.27 |
| hip_flex | 0.96 | 0.10 | 0.99 | 0.01 | 0.95 | 0.14 | 0.97 | 0.04 | 0.66 | 0.32 | 0.88 | 0.12 |
| hip_ab | 0.89* | 0.15 | 0.96* | 0.02 | 0.69 | 0.35 | 0.76 | 0.28 | 0.19 | 0.20 | 0.31 | 0.20 |
| hip_rot | 0.94 | 0.13 | 0.97 | 0.01 | 0.90 | 0.20 | 0.96 | 0.05 | 0.55 | 0.36 | 0.76 | 0.17 |
| lumb_flex | 0.71 | 0.18 | 0.69 | 0.24 | 0.46 | 0.28 | 0.56 | 0.31 | 0.01 | 0.17 | 0.22 | 0.28 |
| lumb_bend | 0.88 | 0.16 | 0.90 | 0.15 | 0.40 | 0.33 | 0.56 | 0.32 | 0.08 | 0.18 | -0.04 | 0.21 |
| lumb_rot | 0.93 | 0.14 | 0.98 | 0.02 | 0.82 | 0.26 | 0.88 | 0.26 | 0.31 | 0.29 | 0.38 | 0.38 |

**Table 5:** Mean (M) and standard deviation (SD) among participants of for trial-to-trial similarity measured by the maximum cross-correlation coefficient $r_{max}$ for all conditions and joint angles. ANOVA revealed significant effects of TASK in all joints Significant differences observed by contrasts are indicated by *.

| | LINE | | | | BEAM | | | | TIGHTROPE | | | |
|---|---|---|---|---|---|---|---|---|---|---|---|---|
| | start | | end | | start | | end | | fail | | succ | |
| | *M* | *SD* | *M* | *SD* | *M* | *SD* | *M* | *SD* | *M* | *SD* | *M* | *SD* |
| ankle_flex | 0.94 | 0.05 | 0.97 | 0.01 | 0.89 | 0.11 | 0.93 | 0.05 | 0.73 | 0.18 | 0.81 | 0.15 |
| knee_flex | 0.97 | 0.06 | 0.98 | 0.01 | 0.96 | 0.08 | 0.97 | 0.04 | 0.95 | 0.04 | 0.96 | 0.03 |
| hip_flex | 0.96 | 0.08 | 0.99 | 0.01 | 0.94 | 0.13 | 0.97 | 0.03 | 0.95 | 0.03 | 0.96 | 0.03 |
| hip_ab | 0.97 | 0.03 | 0.99 | 0.01 | 0.92 | 0.07 | 0.90 | 0.13 | 0.56 | 0.19 | 0.66 | 0.14 |
| hip_rot | 0.90 | 0.13 | 0.96 | 0.04 | 0.87 | 0.14 | 0.93 | 0.04 | 0.78 | 0.19 | 0.86 | 0.17 |
| lumb_flex | 0.83 | 0.20 | 0.90 | 0.17 | 0.79* | 0.19 | 0.91* | 0.14 | 0.61* | 0.17 | 0.80* | 0.16 |
| lumb_bend | 0.85 | 0.08 | 0.88 | 0.15 | 0.65* | 0.18 | 0.68* | 0.19 | 0.49 | 0.09 | 0.49 | 0.13 |
| lumb_rot | 0.92 | 0.07 | 0.95 | 0.03 | 0.82 | 0.16 | 0.88 | 0.15 | 0.50 | 0.20 | 0.65 | 0.21 |

**Table 6:** Mean (M) and standard deviation (SD) among participants of for trial-to-trial similarity measured by the lag time lag% at the maximum cross-correlation coefficient for all conditions and joint angles. ANOVA revealed significant effects of TASK in hip_rot, hip_ad, lumb_bend, lumb_flex. Significant differences observed by contrasts are indicated by *.

| | LINE | | | | BEAM | | | | TIGHTROPE | | | |
|---|---|---|---|---|---|---|---|---|---|---|---|---|
| | start | | end | | start | | end | | fail | | succ | |
| | *M* | *SD* | *M* | *SD* | *M* | *SD* | *M* | *SD* | *M* | *SD* | *M* | *SD* |
| ankle_flex | 0.22 | 0.44 | 0.00 | 0.00 | 4.35 | 12.20 | 1.04 | 2.85 | 8.75 | 18.91 | 3.18 | 8.21 |
| knee_flex | 0.00 | 0.00 | 0.00 | 0.00 | 0.00 | 0.00 | 0.00 | 0.00 | 0.03 | 0.09 | 0.00 | 0.00 |
| hip_flex | 2.76 | 8.73 | 0.00 | 0.00 | 2.00 | 6.32 | 0.00 | 0.00 | 0.89 | 1.86 | 0.23 | 0.74 |
| hip_ab | 2.42* | 4.79 | 0.55* | 0.56 | 7.51 | 9.51 | 7.75 | 11.83 | 32.02 | 15.47 | 21.63 | 10.85 |
| hip_rot | 3.82 | 9.94 | 0.22 | 0.40 | 4.68 | 12.77 | 0.86 | 1.17 | 13.14 | 13.23 | 3.30 | 8.99 |
| lumb_flex | 10.92 | 15.24 | 4.03 | 10.93 | 17.89* | 17.68 | 4.44* | 11.05 | 24.66* | 16.25 | 11.48* | 11.94 |
| lumb_bend | 2.62 | 5.32 | 5.71 | 14.86 | 19.90 | 14.77 | 19.23 | 17.12 | 40.14 | 8.57 | 44.70 | 12.60 |
| lumb_rot | 2.55 | 6.77 | 0.49 | 1.05 | 8.85 | 13.36 | 4.74 | 11.61 | 31.68 | 18.03 | 18.96 | 14.74 |
